# CTCF Promotes Long-range Enhancer-promoter Interactions and Lineage-specific Gene Expression in Mammalian Cells

**DOI:** 10.1101/2020.03.21.001693

**Authors:** Naoki Kubo, Haruhiko Ishii, Xiong Xiong, Simona Bianco, Franz Meitinger, Rong Hu, James D. Hocker, Mattia Conte, David Gorkin, Miao Yu, Bin Li, Jesse R. Dixon, Ming Hu, Mario Nicodemi, Huimin Zhao, Bing Ren

## Abstract

Topologically associating domains (TAD) and insulated neighborhoods (INs) have been proposed to constrain enhancer-promoter communications to enable cell-type specific transcription programs, but recent studies show that disruption of TADs and INs resulted in relatively mild changes in gene expression profiles. To better understand the role of chromatin architecture in dynamic enhancer-promoter contacts and lineage-specific gene expression, we have utilized the auxin-inducible degron system to acutely deplete CTCF, a key factor involved in TADs and IN formation, in mouse embryonic stem cells (mESCs) and examined chromatin architecture and gene regulation during neural differentiation. We find that while CTCF depletion leads to global weakening of TAD boundaries and loss of INs, only a minor fraction of enhancer-promoter contacts are lost, affecting a small subset of genes. The CTCF-dependent enhancer-promoter contacts tend to be long-range, spanning hundreds of kilobases, and are established directly by CTCF binding to promoters. Disruption of CTCF binding at the promoter reduces enhancer-promoter contacts and transcription, while artificial tethering of CTCF to the promoter restores the enhancer-promoter contacts and gene activation. Genome-wide analysis of CTCF binding and gene expression across multiple mouse tissues suggests that CTCF-dependent promoter-enhancer contacts may regulate expression of additional mouse genes, particularly those expressed in the brain. Our results uncover both CTCF-dependent and independent enhancer-promoter contacts, and highlight a distinct role for CTCF in promoting enhancer-promoter contacts and gene activation in addition to its insulator function.

## Introduction

Transcriptional regulation in mammalian cells is orchestrated by cis-regulatory elements that include promoters, enhancers, insulators and other less well characterized sequences^1, 2^. Large-scale projects such as ENCODE have annotated millions of candidate cis-regulatory elements in the human genome and genomes of other mammalian species^3–5^. A majority of these candidate regulatory elements are located far from transcription start sites(i.e. promoters), display tissue and cell-type specific chromatin accessibility, and likely act as enhancers to regulate cell-type specific gene expression. Enhancers are frequently found to be positioned close to their target gene promoters in 3D space at the time of gene activation, suggesting a role for the chromatin architecture in gene regulation ^6, 7^. Indeed, artificially induced spatial proximity between enhancers and promoters has been shown to lead to gene activation ^8, 9^. Insulators, on the other hand, act to block enhancer-promoter contacts to prevent ectopic gene activation ^10–12^. Clearly, in-depth knowledge of the chromatin architecture in each cell type and developmental stage is necessary for mechanistic understanding and functional annotation of enhancers and insulators in the genome.

In recent years, great strides have been made in our understanding of chromatin architecture, thanks to the development of high throughput technologies to capture chromosome conformation^13–19^. These studies have shown that interphase chromosomes reside in separate nuclear space known as chromosome territories, and each chromosome is further partitioned into topologically associating domains (TADs) characterized by higher levels of interactions among DNA within each domain than between domains^1–5^. Within TADs, genes and their regulatory elements are organized into insulated neighborhoods (INs) formed by CTCF-anchored chromatin loops^17, 20^. Both TADs and INs have been proposed to play important roles in gene regulation by constraining enhancer-promoter contacts^13–17, 20^. Supporting this model, previous studies have shown that deletion, duplication or inversion of TAD boundaries result in dysregulation of gene expression and developmental disorders^7, 21–24^. Mechanistically, TADs and INs are proposed to be formed through cohesin/CTCF mediated loop extrusion. In this model, the cohesin complex moves bidirectionally along the chromatin fiber, and the movement is temporarily arrested by DNA-bound CTCF proteins resulting extrusion of the chromatin segments between the two convergent CTCF binding sites^25, 26^. Consistent with this model, acute depletion of CTCF and cohesin complex leads to global loss of TADs and INs. However, the severe disruption of genome structure only affects expression of a small number of genes, raising questions about the general roles of TAD and INs in gene regulation^27, 28^. In addition, a recent study involving high throughput enhancer perturbation followed by single cell RNA-seq analysis found that functional enhancers predominantly reside very close (within 20 kb) to their target genes^29^. These recent studies raise fundamental questions about the prevalence of long-range acting enhancers and the role of TAD/INs in gene regulation.

Here we use two genome-wide chromatin conformation capture assays, namely in situ Hi-C and PLAC-seq (also known as HiChIP)^30, 31^, to determine the chromatin architecture and enhancer-promoter contacts at high-resolution during neural differentiation of mouse embryonic stem cell (mESC) with continuous depletion of CTCF by auxin-inducible degron^32–34^ (Supplementary Table 1). We found that most enhancer-promoter contacts, especially those at close genomic distances, remained intact upon CTCF loss during cell differentiation despite the global weakening of TAD boundaries. However, we also observed lost and newly formed enhancer-promoter contacts at hundreds of dysregulated genes. We characterized the features of CTCF-dependent genes, and found that their promoters are enriched for CTCF binding sites (CBSs) and devoid of nearby enhancers. We showed that CTCF can directly establish enhancer-promoter contacts at these genes since deletion of CTCF-binding site at the promoter reduces enhancer-promoter contact and gene expression, while artificial tethering of CTCF to the promoter could promote enhancer-promoter contacts and gene activation. Furthermore, we found over 2,300 genes that display a significant correlation between CTCF occupancy at the promoter and tissue-specific gene expression patterns, suggesting a role for CTCF binding in their regulation. Our findings uncovered both CTCF-independent and CTCF-dependent mechanisms of enhancer-promoter communications, and provided evidence for a key role for CTCF in directly promoting enhancer-promoter contacts that are distinct from its function at insulator sequences.

## Results

### CTCF loss leads to weakening of TADs/INs without massive gene dysregulation during mESC differentiation

To investigate the functional role of CTCF in chromatin architecture and gene regulation, we utilized an auxin-inducible degron system to acutely deplete CTCF protein in mESC and examined the impact of CTCF loss on dynamics of gene expression and chromatin architecture during mESC differentiation to neural precursor cells (NPCs) (Fig. 1a). The depletion of CTCF was verified by Western blotting and chromatin occupancy of CTCF was nearly completely lost in both ESCs and NPCs, along with loss of cohesin accumulation, as shown by ChIP-seq analysis (Supplementary Fig. 1 and Supplementary Table 2). The CTCF-depleted cells exhibited a delay in the formation of neuronal axons during neural differentiation treatment with cell colonies remaining in round-shape (Fig. 1b). To determine the impact of CTCF loss on chromatin architecture, we first performed Hi-C analyses in mES cells before and during differentiation to NPC, in the presence and absence of CTCF. TADs are characterized by strong intra-domain interactions and relatively weak inter-domain interactions in Hi-C, and the strength of the TAD boundaries can be defined by the insulation score, a ratio between the number of cross-border interactions and the sum of intra-domain interactions within the two adjacent TADs^35^. As shown in Supplementary Fig. 2, CTCF depletion resulted in global loss of chromatin loops and contacts between convergent CTCF binding sites (genomic distance > 100 kb), supporting an essential role for CTCF in the formation of these chromatin organizational features (Supplementary Fig. 2a). We also observed significant weakening of TAD boundaries and a dramatic loss of INs in both ESC and NPC (Supplementary Fig. 2b–e and Supplementary Table 3). This result is generally consistent with previous findings indicating CTCF’s role in the formation of most TADs in mammalian cells^28^ (Supplementary Fig. 2f–h).

**Fig. 1.**
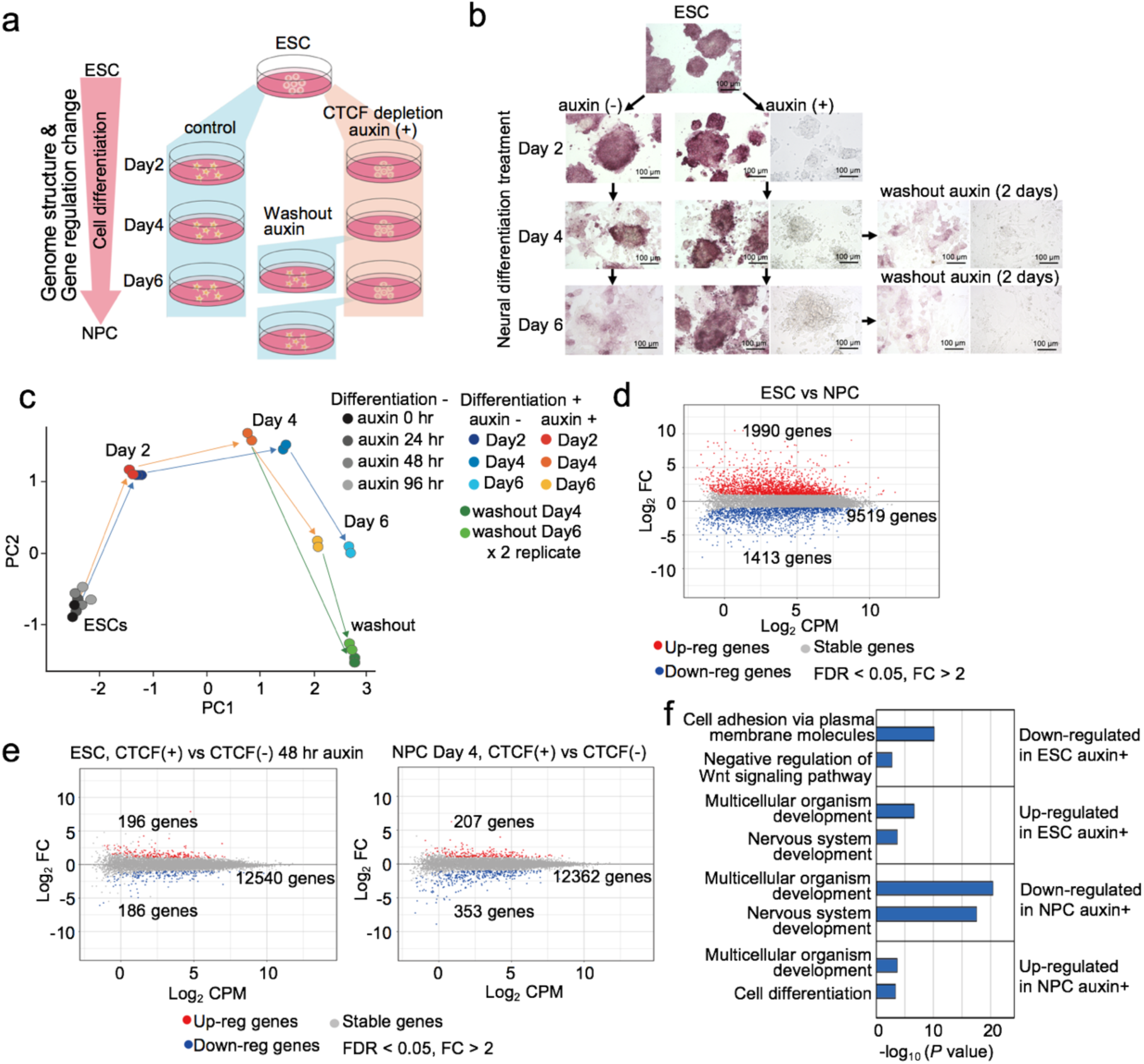
CTCF loss affects only a small fraction of genes in gene regulation during embryonic stem cell differentiation. **a,** Schematic representation of experimental design and sample preparation. Auxin-inducible degron system was utilized to deplete CTCF during cell differentiation from mouse embryonic stem cells (mESCs) to neural progenitor cells (NPCs) (day 2, 4, 6). An additional 2 days of neural differentiation treatment was performed after washing out auxin in day 4 and day 6 differentiated cells. RNA-seq and in situ Hi-C were performed at each time point to examine the impact of CTCF loss on gene regulation and chromatin architecture. **b,** Microscopic images of cell differentiation from mESCs (top) to NPCs (day 2, 4, 6) with or without auxin treatment and additional 2 days of differentiation treatment after washing out auxin from day 4, 6 samples. Alkaline phosphatase staining was performed at every time point. Non-stained bright-field images of each auxin treated sample and auxin washout sample are also shown on the right. **c,** Principal component analysis of gene expression profiles of control and CTCF-depleted cells at each time point of cell differentiation and 2 days after washing out auxin. Gene expression profiles in ESCs with multiple days of auxin treatment (24, 48, and 96 hours) were also analyzed. Two replicates of each sample are shown. **d,** Gene expression changes between control ESCs and NPCs (day 4). Differentially up-regulated and down-regulated genes are plotted in red and blue, respectively (fold change > 2, FDR < 0.05). **e,** Gene expression changes upon CTCF depletion in ESCs (left, 48 hours with or without auxin) and in differentiated cells (right, differentiation day 4 with or without auxin). Differentially up-regulated and down-regulated genes are plotted in red and blue, respectively (fold change > 2, FDR < 0.05). **f,** Top two enriched GO terms of gene sets of differentially expressed genes upon CTCF loss are shown with their p values.

We next investigated the impact of CTCF loss on gene regulation. Consistent with previous reports^28^, the vast majority of genes were expressed normally in CTCF-depleted mESC. Additionally, the gene expression profiles were largely uninterrupted during cell differentiation (Fig. 1c). Only a small fraction of genes (382 genes, 3.0% in ESCs, 560 genes, 4.5% in NPCs) were affected significantly due to CTCF loss (FDR < 0.05, fold change > 2) (Fig. 1d, e, Supplementary Fig. 3, and Supplementary Table 4). Interestingly, genes that are related to neural differentiation (e.g. *Pcdh* cluster genes, *Neurog*, *Neurod4*) were enriched in these CTCF-dependent genes (Fig. 1f), consistent with the observation that CTCF loss is accompanied by abnormal neural differentiation in mESC. Thus, despite the severe disruption of TADs, INs and CTCF-mediated chromatin loops, the gene regulatory programs in mESC during differentiation appear to be mostly unaffected. This observation raised an important question regarding the role of chromatin domains and loops in gene regulation.

### A small subset of enhancer-promoter contacts is dependent on CTCF

To further delineate the relationships between the chromatin structures and gene regulation, we performed quantitative analysis of enhancer-promoter contacts using PLAC-seq (also known as HiChIP)^30, 31^, which interrogates chromatin contacts at select genomic regions at high resolution by combining Hi-C and chromatin immunoprecipitation. We used antibodies against the histone modification H3K4me3, which marks active or poised promoters, to detect chromatin contacts centered on these genomic regions (Fig 2a). We obtained between 300 and 400 million paired-end reads for each replicate (Supplementary Table 1). To determine the differential chromatin contacts in ESCs and NPCs, we analyzed 11,900 gene promoters with similar levels of H3K4me3 ChIP-seq signal using a negative binomial model for each distance-stratified 10-kb interval (Supplementary Fig. 4, Methods). In total, we found 5,913 chromatin contacts between the promoters of 4,573 genes and distal elements to be significantly induced during the neural differentiation (FDR < 0.05), and 1,594 contacts centered on 1,294 genes significantly decreased (Fig. 2b, c). Notably, over 50% of these differential contacts span less than 50 kb in genomic distance (Supplementary Fig. 5a). As expected, these dynamic changes of enhancer-promoter contacts were positively correlated with the changes of active histone modifications such as H3K27ac and H3K4me1 (Supplementary Fig. 5b, c). We confirmed previously reported dynamic enhancer-promoter contacts during mESC cell differentiation (e.g. *Sox2, Hoxb* cluster genes, *Dnmt3b*)^14, 36^ (Fig. 2b, Supplementary Fig. 5d–g).

**Fig. 2.**
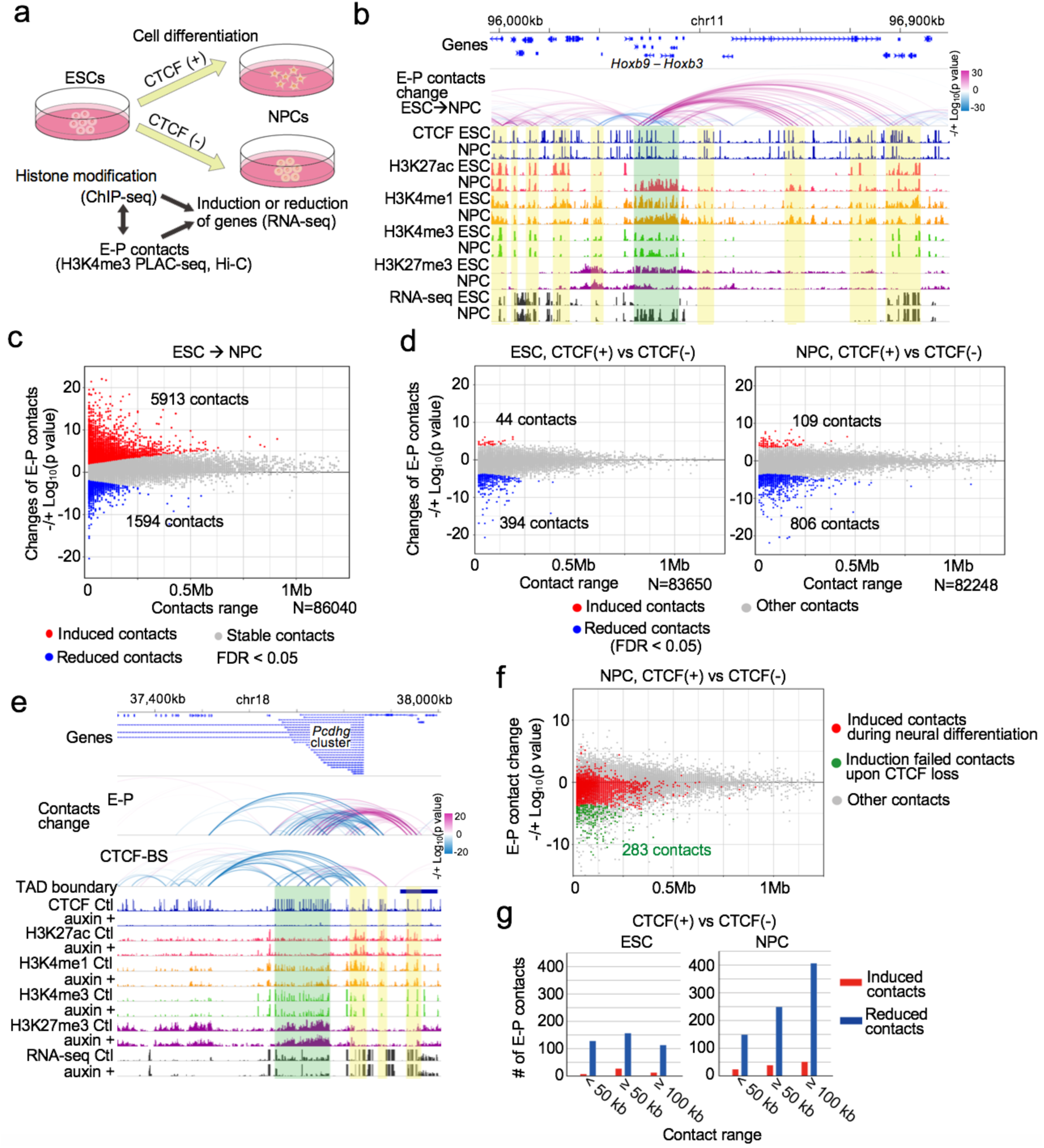
A small number of enhancer-promoter contacts are affected upon CTCF loss despite the drastic weakening of TADs. **a,**Schematic representation of study design to explore the changes of enhancer-promoter (E-P) contacts during neural differentiation in the presence and absence of CTCF. H3K4me3 PLAC-seq datasets were analyzed for changes of E-P contacts at high resolution. ChIP-seq (histone modifications) and RNA-seq (gene regulation) datasets at each time point were also used for combinational analysis. **b,** Genome browser snapshots of a region around *Hoxb3*–*9* genes that were up-regulated during neural differentiation. The arcs show the changes of H3K4me3 PLAC-seq contacts on active elements and promoters identified in differential interaction analysis between ESCs and NPCs (see Methods). The colors of arcs represent degrees of interaction change between compared samples (blue to red, -/+log_10_(p-value)). The promoter regions of *Hoxb3*–*9* genes and interacting enhancer regions are shown in green and yellow shadows, respectively. CTCF, H3K4me1, H3K27ac, H3K4me3, H3K27me3 ChIP-seq and RNA-seq in ESCs and NPCs (day 4) are also shown. **c,** Scatter plots showing genome-wide changes of chromatin contacts anchored on promoters and enhancers (y-axis) identified in differential interaction analysis between ESCs and NPCs. Genomic distances between their two loop anchor sites are plotted in *x*-axis. Significantly induced and reduced chromatin contacts are shown as red and blue dots, respectively (FDR < 0.05). The interaction changes are shown by significance value (-/+log_10_(p-value)) (see Methods). **d,** Scatter plots showing genome-wide changes of chromatin contacts anchored on promoters and enhancers (*y*-axis) identified in differential interaction analysis between control and CTCF-depleted cells in ESC (left) and NPC stage (day 4) (right). Genomic distances between their two loop anchor sites are plotted in *x*-axis. Significantly induced and reduced chromatin contacts are shown as red and blue dots, respectively (FDR < 0.05). **e,** Genome browser snapshots of a region around the *Pcdhg* gene cluster down-regulated upon CTCF loss in NPCs. The arcs show the changes of chromatin contacts on enhancers and promoters (E-P) and chromatin contacts on CTCF binding sites (CTCF-BS) identified in differential interaction analysis between conditions. The colors of arcs represent degrees of interaction change from control cells to CTCF-depleted cells (blue to red, -/+log_10_(p-value)). The promoter regions of *Pcdhg* gene cluster and interacting enhancer regions are shown in green and yellow shadows, respectively. CTCF, H3K4me1, H3K27ac, H3K4me3, H3K27me3 ChIP-seq, and RNA-seq in control and CTCF-depleted NPCs, and TAD boundaries in control cells are also shown. **f,** On the scatter plots of changes of E-P contacts upon CTCF loss in NPCs (Fig. 2d, right panel), E-P contacts that were significantly induced during neural differentiation were plotted in red and significantly reduced contacts upon CTCF loss among them were plotted in green. **g,** Histograms showing the number of significantly changed E-P contacts upon CTCF loss and their genomic distances in ESC (left) and NPC (right) stages. Significantly induced and reduced contacts are shown as red and blue bars, respectively.

Using the same approach, we determined the chromatin contacts dependent on CTCF in mESCs and NPCs. The chromatin contacts between convergent CTCF-binding sites were severely reduced upon CTCF loss (Supplementary Fig. 7a, b), consistent with the results from Hi-C assays. However, the majority of chromatin contacts between enhancers and promoters were unchanged despite the global weakening of TADs and disruption of chromatin loops and INs. Chromatin contacts between 394 and 806 enhancer-promoter pairs in mESCs and NPCs, respectively, decreased significantly upon CTCF loss (FDR < 0.05), while chromatin contacts between 44 and 109 enhancer-promoter pairs in mESCs and NPCs, respectively, increased upon CTCF loss (Fig. 2d, e, Supplementary Table 5). Only 283 pairs of enhancer-promoter contacts out of 5,913 that are normally induced during differentiation failed to be induced in the absence of CTCF (Fig. 2f). Interestingly, genomic distances of the CTCF-dependent enhancer-promoter contacts in differentiated NPCs are generally longer than that in undifferentiated ESCs (Fig. 2g). The modest changes in enhancer-promoter contacts upon CTCF loss are consistent with the mild changes in gene expression in these cells.

### CTCF directly promotes enhancer-promoter contacts and gene expression through binding to gene promoters

We next investigated the features of CTCF-dependent/-independent enhancer-promoter contacts and gene regulation. Since ChIP-seq levels of histone modifications (H3K27ac, H3K4me1, and H3K4me3) were virtually unaffected by CTCF depletion, the observed changes in enhancer-promoter contacts in mESC and NPC were likely a direct consequence of CTCF loss (Supplementary Fig. 4a, Supplementary Fig. 6). Consistent with this interpretation, the anchors of CTCF-dependent enhancer-promoter contacts were strongly enriched for the CBSs detected by ChIP-seq in the mESC and NPC (Fig. 3a). Furthermore, the degree of this enrichment increased with the number of CBSs around the anchors (Fig. 3b). Many such enhancer-promoter contacts that were close to CBSs at their anchor sites were identified in genes that were down-regulated upon CTCF loss (Fig. 3c, d, Supplementary Fig. 7c). In these CTCF-dependent down-regulated genes, we also observed many reduced enhancer-promoter contacts that had CBSs at only one side of their anchor sites, preferentially at promoter side (Fig. 3c, d, Supplementary Fig. 7d, e). These findings indicate that CTCF directly modulates transcription of select genes by binding to their promoters and promoting long-range enhancer-promoter contacts. By contrast, the anchors of enhancer-promoter contacts gained upon CTCF loss were not enriched for CBS, and they were likely a consequence of loss of insulation due to the weakened TAD boundaries and INs (Supplementary Fig. 7f–h).

**Fig. 3.**
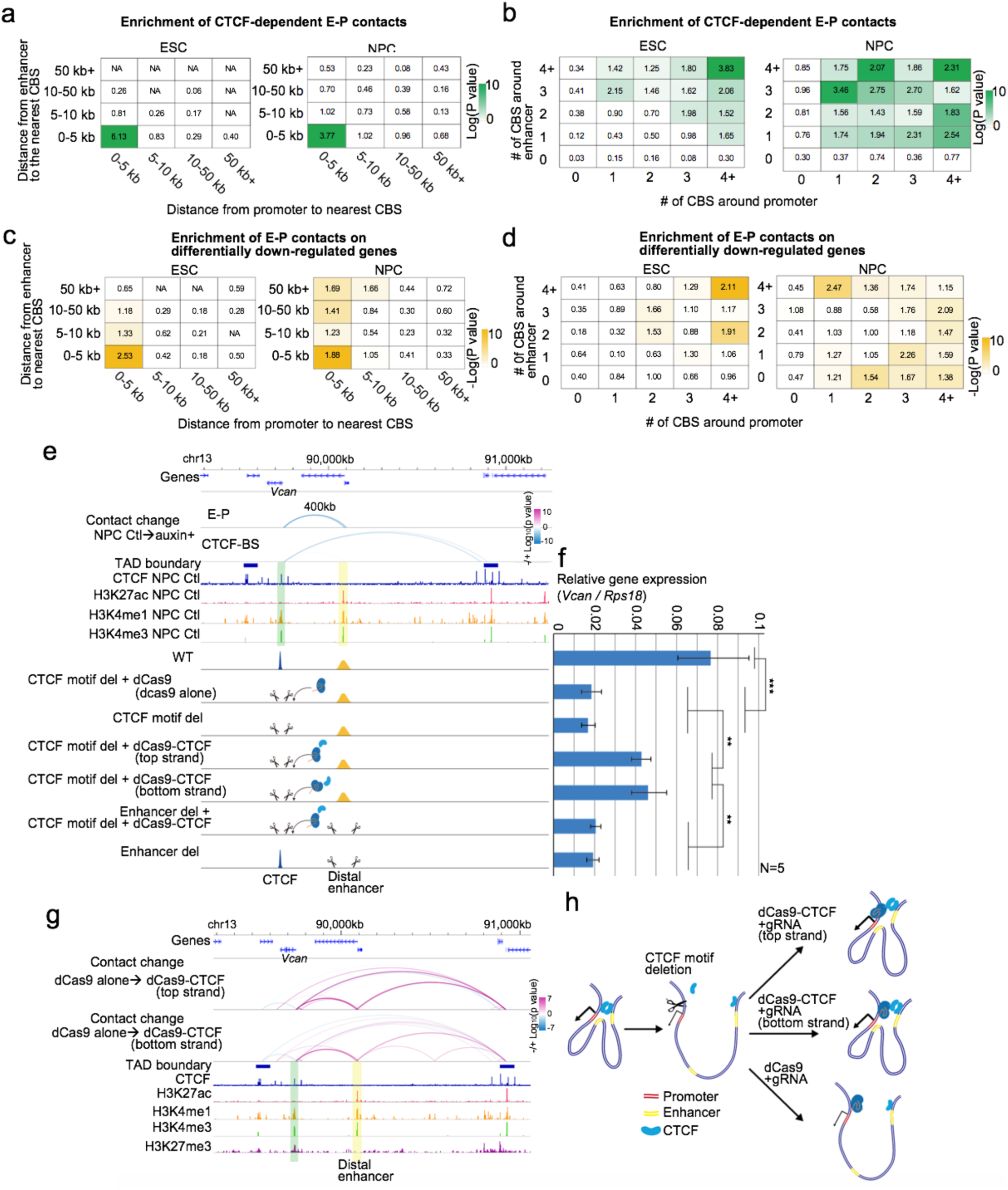
Enriched CBSs at promoters of CTCF-dependent genes and rescue of CTCF-dependent enhancer-promoter contacts and gene regulation by artificially tethered CTCF. **a, b,** Enrichment analysis of CTCF-dependent E-P contacts categorized based on the distance from the loop anchor sites on enhancer side (vertical columns) or promoter side (horizontal columns) to the nearest CTCF binding site (CBS) (a). The same enrichment analysis categorized based on the number of CBSs around loop anchor sites (10 kb bin ±5 kb) on enhancer side (vertical columns) or promoter side (horizontal columns) was also performed (b). Enrichment values are shown by odds ratio (scores in boxes) and p-values (color) in ESC (left) and NPC stage (right) (see Methods). **c, d,** Enrichment analysis of E-P contacts on CTCF-dependent down-regulated genes categorized based on the distance from the loop anchor sites on enhancer side (vertical columns) or promoter side (horizontal columns) to the nearest CBS (c). The same enrichment analysis categorized based on the number of CBSs around loop anchor sites (10 kb bin ±5 kb) on enhancer side (vertical columns) or promoter side (horizontal columns) was also performed (d). Enrichment values are shown by odds ratio (scores in boxes) and p-values (color) in ESC (left) and NPC stage (right) (see Methods). **e,** Genome browser snapshots of a region around *Vcan* gene down-regulated upon CTCF loss in NPCs. The arcs show the changes of chromatin contacts on enhancers and promoters (E-P) and chromatin contacts on CTCF binding sites (CTCF-BS) identified in differential interaction analysis between conditions. The colors of arcs represent degrees of interaction change from control to CTCF-depleted cells (blue to red, -/+log_10_(p-value)). *Vcan* promoter and 400 kb downstream distal active element are shown in green and yellow shadows, respectively. CTCF, H3K4me1, H3K27ac, H3K4me3 ChIP-seq and TAD boundaries in control NPCs are also shown. Schematic representation and description of each cell line is aligned on the bottom. **f,** RT-qPCR expression levels of *Vcan* gene in each cell line in NPC stage. (** p value < 0.01 and *** p value < 0.001, two-tailed t-test). **g,** Changes of chromatin contacts around *Vcan* gene upon artificial tethering of CTCF at the promoter on top and bottom strand are shown. The arcs show the changes of chromatin contacts anchored on enhancers, promoters, and CTCF binding sites identified in differential interaction analysis between conditions. The colors of arcs represent degrees of interaction change from control NPCs (dCas9 alone) to dCas9-CTCF tethered NPCs (blue to red, -/+log_10_(p-value)). CTCF, H3K4me1, H3K27ac, H3K4me3, H3K27me3 ChIP-seq and TAD boundaries in NPCs are also shown. **h,** Schematic representation of observed findings in the rescue experiments. The deletion of the CTCF binding motif adjacent to *Vcan* promoter disrupts its E-P contact along with loss of the CTCF-anchored loop (Supplementary Fig. 9a). Then, the artificially tethered CTCF at the promoter on top or bottom strand rescues the E-P contact and gene regulation.

To confirm that CTCF directly mediates enhancer-promoter contacts to modulate gene expression, we used CRISPR-mediated genome editing to delete a 118-bp sequence containing the CTCF binding motif at the promoter of *Vcan* gene, which encodes a protein that plays an important role in axonal outgrowth^37^ and neural differentiation^38^. *Vcan* is induced during NPC differentiation, and the induction is lost upon CTCF depletion along with a long-range enhancer-promoter contact (400 kb range) anchored by a CBS only on the promoter side. Polymer modelling based on the strings and binders switch (SBS) model also supports such changes of chromatin contacts upon CTCF depletion^39, 40^ (Supplementary Fig. 8). Upon removal of the CTCF binding sequence, *Vcan* expression was significantly reduced in NPC cells. This reduction in *Vcan* expression could be largely restored by tethering the CTCF protein to the mutated *Vcan* promoter using a dCas9-CTCF fusion and a guide RNA (gRNA) targeting a sequence adjacent to the deleted CTCF binding sequence, in two different experiments using distinct gRNAs (Fig. 3e, f, Supplementary Fig. 9). The rescue of the *Vcan* expression by the artificially tethered CTCF was dependent on the distal element sequences (Fig. 3e, f). PLAC-seq experiments showed that the artificially tethered CTCF protein could re-establish the long-range enhancer-promoter contact at *Vcan* that was lost upon deletion of the CTCF binding motif near the promoter (Fig. 3g, h). Taken together, the above results demonstrated that CTCF can directly promote long-range enhancer-promoter contact and gene regulation by binding to the gene promoter.

### CTCF-dependent genes reside in enhancer deserts

While many of the genes with CBSs at promoters were dependent on CTCF in mESCs and NPCs, many others were not affected by CTCF loss despite having CTCF bindings at gene promoters (Supplementary Fig. 10a). These CTCF-independent genes are generally close to the genomic regions associated with the H3K27ac histone mark (≤ 50 kb, PLAC-seq peak signal p-value < 0.01) (Fig. 4a, b, Supplementary Fig. 10b, c), implying that they are regulated by short-range enhancer-promoter contacts formation of which are independent of CTCF (Fig. 4b, Supplementary Fig. 10d, e). By contrast, CTCF-dependent genes were generally regulated by long-range enhancer-promoter contacts (≥ 100 kb, Fig. 4c) especially in NPCs (Fig. 4c, Supplementary Fig. 10f). Similarly, genes up-regulated upon CTCF depletion differed from those down-regulated in whether they were located at enhancer desert regions or not. While the down-regulated genes tended to be located at enhancer desert regions (2 enhancers or less around transcription start site (TSS) < 200 kb in Fig. 4c), the up-regulated genes were close to multiple enhancers (Fig. 4c, d, and Supplementary Fig. 11 for their examples).

**Fig. 4.**
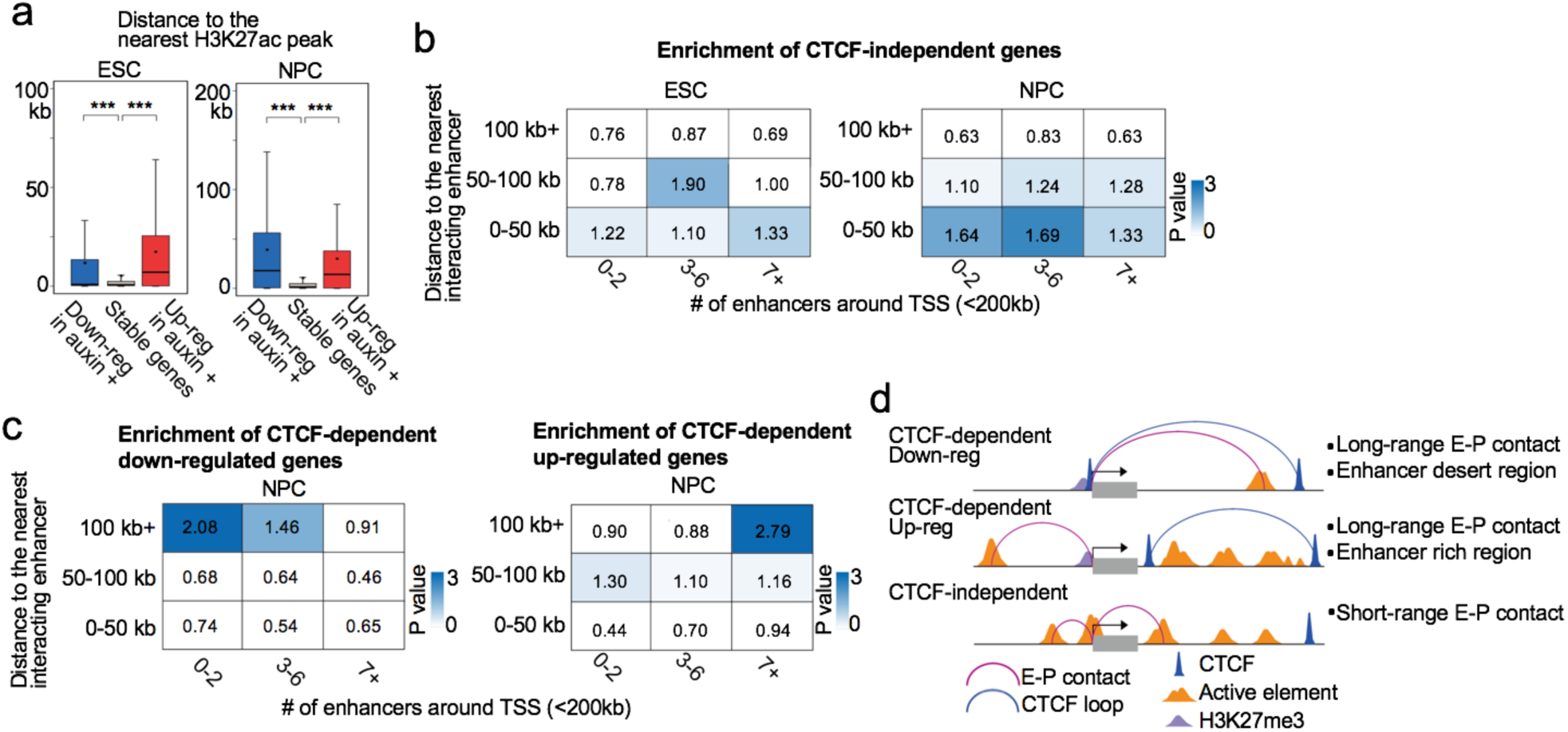
General features of CTCF-dependent/-independent genes. **a,**Boxplots showing the distance from transcription start site (TSS) to the nearest H3K27ac peak in ESC (left) and NPC stage (right). Red: CTCF-dependent up-regulated genes, blue: CTCF-dependent down-regulated genes, gray: CTCF-independent stably regulated genes. (*** p value < 0.001, two-tailed t-test). **b,** Enrichment analysis of CTCF-independent genes categorized based on the distance to the nearest interacting enhancer (vertical columns) and the number of enhancers around TSS (< 200 kb) (horizontal columns) in ESCs (left) and NPCs (right). Enrichment values are shown by odds ratio (scores in boxes) and p-values (color). The distance to the nearest interacting enhancer is represented by the shortest genomic distance of significant PLAC-seq peaks on enhancers and promoters (p-value < 0.01) (see Methods). **c,** Enrichment analysis of CTCF-dependent down-regulated genes (left) and up-regulated genes (right) categorized based on the distance to the nearest interacting enhancer (vertical columns) and the number of enhancers around TSS (< 200 kb) (horizontal columns) in NPCs. Enrichment values are shown by odds ratio (scores in boxes) and p-values (color). The distance to the nearest interacting enhancer is represented by the shortest genomic distance of significant PLAC-seq peaks on enhancers and promoters (p-value < 0.01) (see Supplementary Fig. 10 f for the same analysis in ESCs and Methods). **d,** Model for the general features of CTCF-dependent down-regulated (top), up-regulated genes (middle), and CTCF-independent stably regulated genes (bottom).

### Promoter occupancy by CTCF correlates with expression at lineage-specific genes across diverse mouse tissues

The above findings suggest a previously under-appreciated mechanism for CTCF in gene regulation. In contrast to its well-established role in forming chromatin loops, TAD boundaries and insulators, we demonstrated that CTCF also directly binds to gene promoters to promote long-range enhancer-promoter contacts and enable enhancer-dependent gene expression in mammalian cells. In mouse ESCs and NPCs, several hundred genes are subject to regulation by this mechanism. These include the proto-cadherin gene clusters that were previously reported to be regulated by CTCF binding sites at the promoters and the distal enhancer^41^ (Fig. 2e). To further explore the extent of genes subject to this CTCF-dependent mechanism, we examined public ChIP-seq datasets of CTCF binding and RNA-Seq across multiple mouse tissues (9 tissue samples from ENCODE^4, 42^, Supplementary Table 6). Consistent with this postulated mechanism, CBSs are enriched around promoters (Fig. 5a, Supplementary Fig. 12a) and ChIP-seq signals around promoters (TSS ±10 kb) show positive correlation with gene expression in over 2,300 mouse genes in these tissues (r > 0.6, 2,332 genes), many of which could not be explained by DNA methylation levels at the promoter-proximal CBSs (Fig. 5b, Supplementary Fig. 12b). Interestingly, high lineage-specificity in transcription as measured by Shannon entropy^43^ was predominantly found in the forebrain-specific genes and the most enriched gene ontology (GO) term in this gene group was related to “synapse assembly”. On the other hand, GO terms related to “signaling pathway” were enriched in the other tissue-specific genes (Fig. 5c, Supplementary Fig. 12c–e). Many forebrain-specific and heart-specific genes were down-regulated in CTCF-depleted NPCs and CTCF knockout heart tissue^44^, respectively (Supplementary Fig. 12f, g), supporting that many of these genes are indeed regulated by CTCF binding to the gene promoters in a lineage-specific manner.

**Fig. 5.**
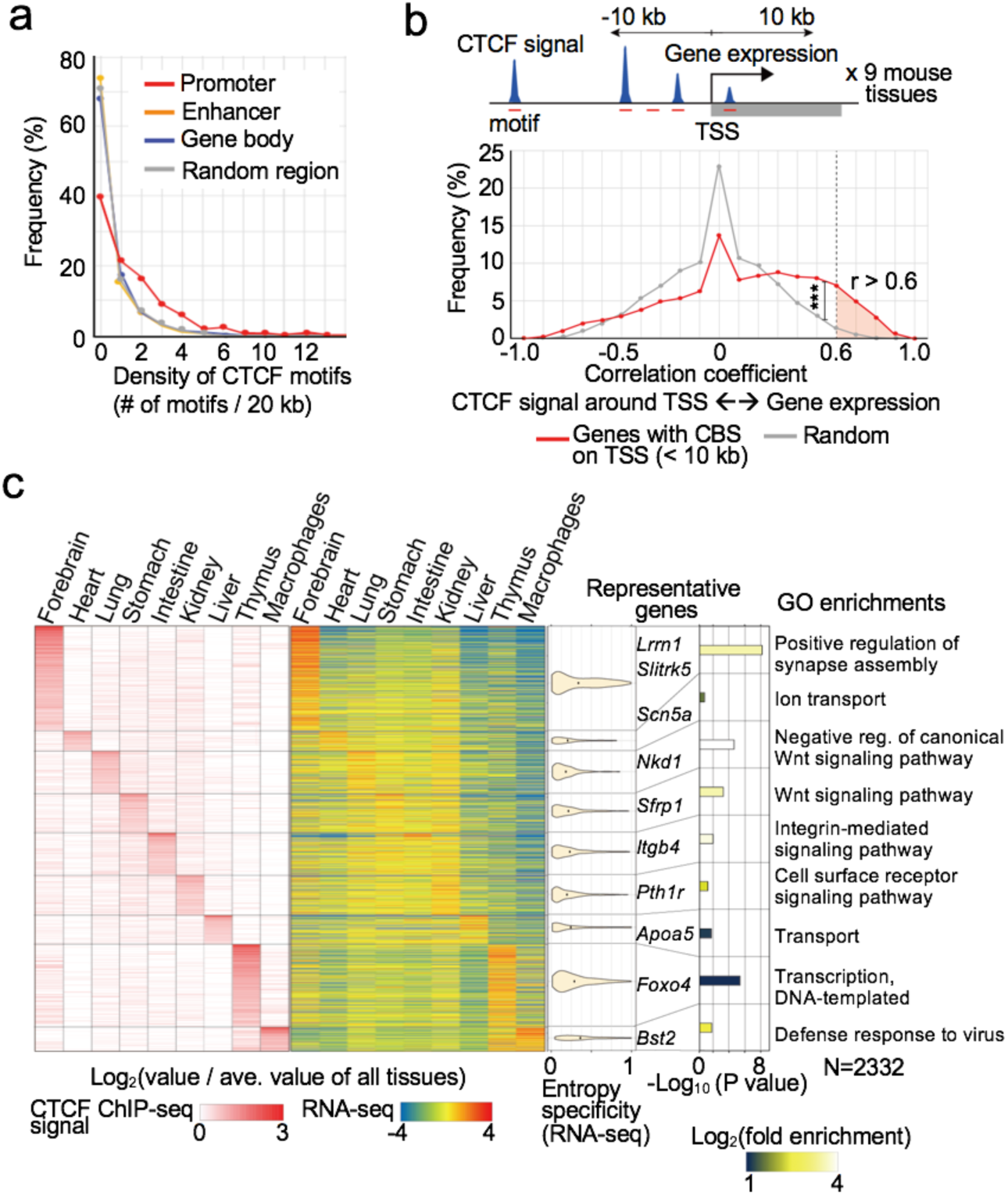
CTCF bindings on gene promoters correlates with tissue-specific gene regulation. **a,**Frequency of genomic regions and their density of CTCF motifs are plotted. The genomic regions were classified into promoters, enhancers (identified in ESCs and NPCs), gene bodies, and random regions. **b,** Schematic representation of data analysis for correlation between the sum of CTCF ChIP-seq signals around TSS (< 10 kb) and gene expression levels across multiple mouse tissues (top, see Methods). Frequencies of genes are plotted based on the correlation coefficient between the CTCF ChIP-seq signals around TSS and gene expression levels across multiple mouse tissues (red line). The same plots analyzed using randomly shuffled CTCF ChIP-seq datasets are shown in gray as control (*** p value < 0.001, Pearson’s Chi-squared test for the comparison of fraction of genes with positive correlation coefficient (r ≥ 0.6) or others (r < 0.6) between the two groups). **c,** Heatmaps showing the lineage-specificity of CTCF ChIP-seq signals around promoters and their gene expression levels. Genes that had high correlation coefficient (> 0.6) in panel (b) are shown (2,332 genes). The lineage-specificity was calculated by log_2_(value / average value of all tissues). The violin plots also show the lineage-specificity of transcription measured by Shannon entropy in each gene group. The width is proportional to the sample size. Top enriched GO terms of each group genes are shown with fold enrichment, p-value, and their representative genes.

## Discussion

CTCF- and cohesin-mediated chromatin structures such as TADs and INs^17, 27, 28^ have been postulated to play a role in constraining enhancer-promoter communications^13–16^. However, the vast majority of genes are expressed normally in the absence of CTCF or Cohesin^27, 28^, raising questions about the role of chromatin architecture, especially enhancer-promoter contacts, in gene regulation. Here, we provided multiple layers of evidence that CTCF not only actively forms TADs and INs, but also directly promotes enhancer-promoter contacts and potentially contributes to activation of thousands of lineage-specific genes. CTCF binding to the promoter of such genes is necessary and sufficient for establishing their enhancer-promoter contacts. We demonstrated that artificial tethering of CTCF to gene promoter could promote enhancer-promoter contacts and gene activation. Further analysis of tissue-specific CTCF binding profiles and gene expression patterns across multiple mouse tissues uncovered several thousand genes that might be regulated by CTCF-dependent promoter-enhancer contacts.

Meanwhile, our study revealed that the majority of enhancer-promoter contacts are independent of CTCF, which could explain the modest change of gene expression profiles upon CTCF loss. Most of the enhancer-promoter contact changes during cell differentiation were associated with the enhancer activities. In addition to the enhancer activity itself, it can be assumed that these CTCF-independent enhancer-promoter contacts are mediated by other factors. For example, Yin Yang 1 (YY1) has been shown to regulate enhancer-promoter contacts in mouse embryonic stem cells^45, 46^. Another genomic interaction mediator, LIM-domain-binding protein 1 (LDB1), is also known to control long-range and trans interactions^47–49^ that regulate specific gene sets such as olfactory receptor genes^49^ and genes for cardiogenesis^48^.

Our study also highlights the biological importance of the long-range chromatin contacts in lineage-specific gene expression. A recent study showed that functional enhancer-promoter pairs predominantly locate in very close genomic distances^29^, despite the fact that a huge number of cis-regulatory elements and interactions between promoters and distal elements have been annotated^1, 2, 50, 51^. We show here that lineage-specific expression of many genes may be dependent on long-range (>100 kb) enhancer-promoter contacts anchored by CTCF binding at the promoters. It is reasonable to assume that some specific long-range enhancer-promoter contacts that potentially determine lineage-specificity require the rigid structure of long-range CTCF loops^45, 46^. Consistent with this model of CTCF function in promoting long-range enhancer-promoter contacts and lineage specific gene expression, many previous studies that have shown that cell-type specific CTCF depletion or deletion in the mouse leads to severe developmental defects^7^.

CTCF has been implicated in a variety of human diseases. It has been previously reported that CBSs are highly mutated in several cancer types^52–54^ and somatic CTCF mutations also occur in about one-quarter of endometrial carcinoma^55^. Thus, further study of the mechanism for CTCF in gene regulation will help to elucidate the role of CTCF and chromatin organization in tumorigenesis and non-coding cancer drivers.

## Methods

### Cell culture

The F1 Mus musculus castaneus × S129/SvJae mouse ES cells (XY; F123 cells)^56^ (a gift from Rudolf Jaenisch) were cultured in KnockOut Serum Replacement containing mouse ES cell media: DMEM 85%, 15% KnockOut Serum Replacement (Gibco), penicillin/streptomycin (Gibco), 1× non-essential amino acids (Gibco), 1× GlutaMax (Gibco), 1000 U/ml LIF (Millipore), 0.4 mM β-mercaptoethanol. The cells were typically grown on 0.2% gelatin-coated plates with irradiated mouse embryonic fibroblasts (MEFs) (GlobalStem). Cells were maintained by passaging using Accutase (Innovative Cell Technologies) on 0.2% gelatin-coated dishes (GENTAUR) at 37°C and 5% CO2. Medium was changed daily when cells were not passaged. Cells were checked for mycoplasma infection and tested negative.

### Construction of the plasmids

The CRISPR/Cas9 plasmid (CTCF-mouse-3sgRNA-CRISPRexp-AID) was assembled using the Multiplex CRISPR/Cas9 Assembly System kit^57^ (a gift from Takashi Yamamoto, Addgene kit #1000000055). Oligonucleotides for three gRNA templates were synthesized, annealed and introduced into the corresponding intermediate vectors. The first gRNA matches the genome sequence 23 bp upstream of the stop codon of mouse CTCF. The oligonucleotides with sequences (5’-CACCGTGATCCTCAGCATGATGGAC-3’) and (5’-AAACGTCCATCATGCTGAGGATCAC-3’) were annealed. The other two gRNAs direct *in vivo* linearization of the donor vector: the first pair of oligonucleotides are (5’-CACCGCTGAGGATCATCTCAGGGGC-3’) and (5’-AAACGCCCCTGAGATGATCCTCAGC-3’); the second pair is (5’-CACCGATGCTGGGGCCTTGCTGGC-3’) and (5’-AAACGCCAGCAAGGCCCCAGCATC-3’). The three gRNA-expressing cassettes were incorporated into one single plasmid using Golden Gate assembly. The donor vector (mCTCF24-AID-donor-Neo) was constructed using PCR and Gibson Assembly Cloning kit (New England Biolabs). The insert cassette includes sequences that codes for a 5GA linker, the auxin-induced degron (AID), a T2A peptide and the neomycin resistant marker, and is flanked by 24-bp homology arms to integrate into the CTCF locus. The left and right arms have sequences CCTGAGATGATCCTCAGCATGATG and GACCGGTGATGCTGGGGCCTTGCT, respectively. The AID coding sequence was amplified from pcDNA5-H2B-AID-EYFP^33^ (a gift from Don Cleveland, Addgene plasmid #47329) and the T2A-Neo^R^ was amplified from pAC95-pmax-dCas9VP160-2A-neo^58^ (a gift from Rudolf Jaenisch, Addgene plasmid #48227). The sequence for the 5GA linker was included in one of the primers. The original donor backbone was a gift from Dr. Ken-ichi T. Suzuki from Hiroshima University, Hiroshima, Japan.

The donor vector encodes the following amino acid sequence that corresponds to the 24-bp left homology arm of CTCF, a 5GA linker, AID, T2A, and Neo^R^: PEMILSMMGAGAGAGAGAGSVELNLRETELCLGLPGGDTVAPVTGNKRGFSETVDLKLNLNNEPANKEGSTTHDVVTFDSKEKSACPKDPAKPPAKAQVVGWPPVRSYRKNVMVSCQKSSGGPEAAAFVKVSMDGAPYLRKIDLRMYKSYDELSNALSNMFSSFTMGKHGGEEGMIDFMNERKLMDLVNSWDYVPSYEDKDGDWMLVGDVPWPMFVDTCKRLRLMKGSDAIGLAPRAMEKCKSRAGSGEGRGSLLTCGDVEENPGPRLETRMGSAIEQDGLHAGSPAAWVERLFGYDWAQQTIGCSDAAVFRLSAQGRPVLFVKTDLSGALNELQDEAARLSWLATTGVPCAAVLDVVTEAGRDWLLLGEVPGQDLLSSHLAPAEKVSIMADAMRRLHTLDPATCPFDHQAKHRIERARTRMEAGLVDQDDLDEEHQGLAPAELFARLKARMPDGEDLVVTHGDACLPNIMVENGRFSGFIDCGRLGVADRYQDIALATRDIAEELGGEWADRFLVLYGIAAPDSQRIAFYRLLDEFF*.

The lentiviral vector for expressing TIR1 (Lentiv4-EFsp-Puro-2A-TIR1-9Myc) was constructed using PCR and Gibson Assembly Cloning kit (New England Biolabs). The backbone was modified from lentiCRISPR v2^59^ (a gift from Feng Zhang, Addgene plasmid #52961) and the TIR1-9myc fragment was amplified from pBabe TIR1-9myc^33^ (a gift from Don Cleveland, Addgene plasmid #47328). The expressing cassette includes a puromycin resistant marker followed by sequences that code for P2A peptide and TIR1-9myc protein. The gene expression is driven by EFS promoter in the original lentiCRISPR v2. The maps and the sequences of the plasmids are available at the following URLs. CTCF-mouse-3sgRNA-CRISPRexp-AID (https://benchling.com/s/seq-1R4nJ8quYptUqerRWSdX), mCTCF24-AID-donor-Neo (https://benchling.com/s/seq-LtJu9OTscKJNCEMOk8ok), Lentiv4-EFsp-Puro-2A-TIR1-9Myc (https://benchling.com/s/seq-6wSCsW3Kr9S1igXZ8H9K).

### Transfection and establishment of CTCF-AID knock-in clones

The cells were passaged once on 0.2% gelatin-coated feeder-free plates before transfection. The cells were transfected using the Mouse ES Cell Nucleofector Kit (Lonza) and Amaxa Nucleofector (Lonza) with 10 µg of the CRISPR plasmid and 5 µg of the donor plasmid following the manufacturer’s instructions. After transfection, the cells were plated on drug-resistant MEFs (GlobalStem). Two days after transfection, drug selection was started by addition of 160 µg/ml G418 (Geneticin, Gibco) to the medium. Drug-resistant colonies were isolated and the clones with AID knock-in on both alleles were found by performing PCR of the genomic DNA using primers specific to sequences flanking the 3’ end of the CTCF coding sequence (AAATGTTAAAGTGGAGGCCTGTGAG and AAGATTTGGGCCGTTTAAACACAGC). The sequence at the CTCF-AID junction on both alleles were checked by sequencing of allele-specific PCR products, which were generated by using either a CTCF-129-specific (CTGACTTGGGCATCACTGCTG) or a CTCF-Cast-specific (GTTTTGTTTCTGTTGACTTAGGCATCACTGTTA) forward primer and a reverse primer in the AID coding sequence (GAGGTTTGGCTGGATCTTTAGGACA). The expression of CTCF-AID fusion protein was confirmed by observing the difference in the molecular weight compared to the control cells by Western blot with anti-CTCF antibody (Millipore, 07-729).

### Lentivirus production and infection

We produced the lentivirus for expressing TIR1-9myc using Lenti-X Packaging Single Shots system (Clontech) and infected the CTCF-AID knock-in mESCs following the manufacturer’s instructions. After infection, the cells were selected by culturing with 1 µg/ml puromycin. Drug-resistant colonies were isolated and expression of TIR1-9myc was confirmed by Western blot using anti-Myc antibody (Santa Cruz, sc-40). Clones expressing high level of TIR1-9myc were used for the subsequent experiments.

### Preparation of CTCF-depleted cells and neural progenitor cell differentiation

The CTCF-AID knock-in mESCs expressing TIR1-9myc were passaged on 0.2% gelatin-coated plates without MEFs. We added 1 ul 500 mM auxin (Abcam, ab146403) per 1 ml medium to deplete CTCF, and changed medium with auxin every 24 hours. Cells were harvested 24, 48 or 96 hours after starting auxin treatment. For NPC differentiation, the CTCF-AID knock-in mESCs were grown on MEFs and passaged on 0.2% gelatin-coated plates without MEFs one day before starting differentiation treatment. The cells were plated sparsely to avoid passaging to new plates during neural differentiation because most of the cells failed to attach to new plates after auxin treatment. On day 0, auxin was added to the CTCF-depleted cell samples, and LIF was deprived from the culture medium 6 hours after adding auxin. From day 1, 5 uM retinoic acid (Sigma, R2625) was added with LIF-deprived medium and auxin was also added continuously to the CTCF-depleted cell samples. Cells were harvested on day 2, day 4 and day 6. To harvest auxin-washout samples, auxin treatment was stopped on day 4 or day 6 and differentiation treatment was continued for another 2 days. Alkaline phosphatase staining was performed on each time point using the AP Staining kit II (Stemgent, 00-0055).

### Antibodies

Antibodies used in this study were rabbit anti-CTCF (Millipore, 07-729, for western blotting), rabbit anti-Histone H3 (abcam, ab1791, for western blotting), rabbit anti-CTCF (Active Motif, 61311, for microChIP-seq), rabbit anti-Rad21 (Santa Cruz, sc-98784, for microChIP-seq), rabbit anti-H3K4me1 (abcam, ab8895, for ChIP-seq), rabbit anti-H3K4me3 (Millipore, 04-745, for ChIP-seq), rabbit anti-H3K27ac (Active Motif, 39133, for ChIP-seq), mouse anti-H3K27me3 (Active Motif, 61017, for ChIP-seq), mouse anti-Myc antibody (Santa Cruz, sc-40, for western blotting), and mouse anti-Cas9 (Cell Signaling, 14697, for western blotting). Goat anti-Rabbit IgG (H+L)-HRP (Bio Rad, 1706515) and Goat anti-Mouse IgG (H+L)-HRP (Invitrogen, 31430) were used as secondary antibody for western blotting.

### Western blotting

Cells were washed with PBS and scraped in cold PBS, and pelleted to be stored at −80°C. Two million cells were resuspended in 100 μL lysis buffer (20 mM Tris-HCl, 150 mM NaCl, 1 mM EDTA, 1mM EGTA, 1% Triton X-100, 1x complete protease inhibitor (Roche)), and sonicated for 10 minutes total ON time with pulses of 15 second ON and OFF, and 40% amplitude using QSONICA 800R (Qsonica). Protein concentration was measured using Pierce BCA Protein Assay Kit (Thermo Fisher). Laemmli Sample Buffer (Bio-Rad) with 355 mM 2-Mercaptoethanol was mixed with 15 μg of each sample and incubated for 5 minutes at 95°C. The samples were run on 4-15% Mini-PROTEAN® TGX™ Precast Gels (Bio-Rad), and transferred onto nitrocellulose membranes at 100 V for 1 hour. The membranes were rinsed with 1x TBST and blocked with 5% dry milk at room temperature for 45 minutes. After washing with TBST, the membranes were incubated with diluted antibody in the blocking buffer overnight at 4°C. After overnight incubating, membranes were washed 4 times 5minutes in 1x TBST at room temperature, and incubated with secondary antibody in blocking buffer at room temperature for 45 minutes. After washing 4 times with TBST, the substrates were detected using Pierce ECL Western Blotting Substrate (Thermo Fisher).

### Cell cycle analysis

Cells were grown in 6-well plates. After dissociation with Accutase (Innovative Cell Technologies), 2-5 million cells were washed with PBS and re-suspended in 300 µl ice-cold PBS. Cells were fixed for a minimum of 24h at 4°C after drop-wise addition of 800 µl ice-cold ethanol. After fixation, cells were pelleted and re-suspended in PBS containing 0.1% Triton X-100, 20 µg/mL Propidium iodide and 50 µg/ml RNase A. Cells were incubated for 30 min at 37°C before subjected to flow cytometry analysis.

### MicroChIP-seq library preparation

MicroChIP-seq experiments for CTCF and Rad21 were performed as described in ENCODE experiments protocols (“Ren Lab ENCODE Chromatin Immunoprecipitation Protocol for MicroChIP” in https://www.encodeproject.org/documents/) with minor modifications. Cells were crosslinked with 1% formaldehyde for 10 minutes and quenched with 125mM glycine. We used 0.5 million cells for microChIP. Chromatin shearing was performed using truChIP Chromatin Shearing Reagent Kit (Covaris) according to the manufacturer’s instructions. Covaris M220 was used for sonication with the following parameters: 10 minutes duration at 10.0% duty factor, 75.0 peak power, 200 cycles per burst at 5-9°C temperature range. The chromatin was diluted with 10 mM Tris-HCl pH 7.5, 140 mM NaCl, 1 mM EDTA, 0.5 mM EGTA, 1% Triton X-100, 10% SDS, 0.1% Sodium Deoxycholate, 1x complete protease inhibitor (Roche), 1 mM PMSF to adjust to 0.21% SDS concentration. We used 8 μL anti-rabbit IgG Dynabeads (Life Technologies) for CTCF or Rad21 antibodies and washed the beads with cold RIPA buffer 1 (10 mM Tris-HCl pH 7.5, 140 mM NaCl, 1 mM EDTA, 0.5 mM EGTA, 1% Triton X-100, 0.1% SDS, 0.1% Sodium Deoxycholate) for 2 times. After washing, 5 μg antibody (anti-CTCF or anti-Rad21) with 95 μL RIPA buffer 1 was added to the beads and incubated on a rotating platform at 4°C for 6 hours. After incubation, beads were washed once with 100 μL cold RIPA buffer 1 and mixed with chromatin followed by overnight incubation on a rotating platform at 4°C. Beads were washed 4 times with 10 mM Tris-HCl pH 7.5, 300 mM NaCl, 1 mM EDTA, 0.5 mM EGTA, 1% Triton X-100, 0.2% SDS, 0.1% Sodium Deoxycholate and washed once with 100 μL cold 1x TE. After removing the TE, 150 μL elution buffer 1 (20 mM Tris-HCl pH 7.5, 50 mM NaCl, 5 mM EDTA, 1% SDS) was added. The input samples were processed in parallel with the ChIP samples. RNase A (final conc. = 0.2 mg/mL) was added and incubated at 37°C for 1 hour with shaking at 1200rpm. Samples were incubated with proteinase K (final conc. = 0.13 mg/mL) at 68°C for 4 hours with shaking at 1200rpm. After removal of beads, the samples were extracted with Phenol: Chloroform: Isoamyl Alcohol (25:24:1) and precipitated with ethanol. To prepare Illumina sequencing libraries, ThruPLEX DNA-seq 12s kit (Rubicon Genomics) was used according to the manufacturer’s instructions. We used 0.5-1.0 ng IP materials and 50 ng input DNA for library preparation, and 11-12 and 5 cycles of PCR were performed respectively. After purification by 1x AMpure Beads (Beckman Coulter), library quality and quantity were estimated with TapeStation (Agilent Technologies) and Qubit (Thermo Fisher Scientific) assays. Libraries were sequenced on HiSeq2500 or HiSeq4000 single end for 50 bp.

### ChIP-seq library preparation

ChIP-seq experiments for each histone mark were performed as described in ENCODE experiment protocols (“Ren Lab ENCODE Chromatin Immunoprecipitation Protocol” in https://www.encodeproject.org/documents/) with minor modifications. Cells were crosslinked with 1% formaldehyde for 10 minutes and quenched with 125mM glycine. We used 1.0 million cells for each ChIP sample. Shearing of chromatin was performed using truChIP Chromatin Shearing Reagent Kit (Covaris) according to the manufacturer’s instructions. Covaris M220 was used for sonication with following parameters: 10 minutes duration at 10.0% duty factor, 75.0 peak power, 200 cycles per burst at 5-9°C temperature range. The concentration of fragmented DNA was diluted to 0.2 μg/μl with 1x TE. For immunoprecipitation, we used 11 μL anti-rabbit or anti-mouse IgG Dynabeads (Life Technologies) and wash them with cold BSA/PBS (0.5 mg / mL bovine serum albumin in 1x phosphate buffered saline) for 3 times. After washing, 3 μg antibody with 147 μL cold BSA/PBS were added to the beads and incubated on a rotating platform at 4°C for 2 hours. After incubation, beads were washed with150 μL cold BSA/PBS for 3 times, and mixed with 100 μL Binding Buffer (1% Triton X-100, 0.1% Sodium Deoxycholate, 1x complete protease inhibitor (Roche)) plus 100 μL 0.2 μg/μl chromatin followed by overnight incubation on a rotating platform at 4°C. Beads were washed 5 times with 50 mM Hepes pH 8.0, 1% NP-40, 1 mM EDTA, 0.70% Sodium Deoxycholate, 0.5 M LiCl, 1x complete protease inhibitor (Roche) and washed once with 150 μL cold 1x TE followed by incubattion at 65°C for 20 minutes in 150 μL ChIP elution buffer (10 mM Tris-HCl pH 8.0, 1 mM EDTA, 1% SDS). The beads were removed and the samples were further incubated at 65°C overnight to reverse crosslinks. The input samples were processed in parallel with the ChIP samples. Samples were incubated with RNase A (final conc. = 0.2 mg/mL) at 37°C for 1 hour, and Proteinase K (final conc. = 0.4 mg/mL) was added and incubated at 55°C for 1 hour. The samples were extracted with phenol: chloroform: isoamyl alcohol (25:24:1) and precipitated with ethanol. We used 3-5 ng of starting IP materials for preparing Illumina sequencing libraries. The End-It DNA End-Repair Kit (Epicentre) was used to repair DNA fragments to blunt ends, and the samples were purified by Qiagen MinElute PCR Purification kit (Qiagen, Cat#28006). A-tailing 3’ end was performed using Klenow Fragment (3’→5’ exo-) (New England Biolabs), and then TruSeq Adapters were ligated by Quick T4 DNA Ligase (New England Biolabs). Size selection using AMpure Beads (Beckman Coulter) was performed to get 300-500bp DNA and PCR amplification (8-10 cycles) were performed. After purification by 1x AMpure Beads (Beckman Coulter), library quality and quantity were estimated with TapeStation (Agilent Technologies) and Qubit (Thermo Fisher Scientific) assays. Libraries were sequenced on HiSeq4000 single end for 50 bp.

### RNA-seq library preparation

Total RNA was extracted from 1–2 million cells using the AllPrep Mini kit (QIAGEN) according to the manufacturer’s instructions and 1 μg of total RNA was used to prepare each RNA-seq library. The libraries were prepared using TruSeq Stranded mRNA Library Prep Kit (Illumina). Library quality and quantity were estimated with TapeStation (Agilent Technologies) and Qubit (Thermo Fisher Scientific) assays. Libraries were sequenced on HiSeq4000 using 50 bp paired-end.

### Hi-C library preparation

*In situ* Hi-C experiments were performed as previously described using the MboI restriction enzyme^16^. Cells were crosslinked with 1% formaldehyde for 10 minutes. Then 25 μl per mL of 2.5M Glycine was added followed by a 5-minute incubation at room temperature and then a 15-minute incubation on ice. The crosslinked pellets were washed with 1 x PBS and incubated with 200ul of lysis buffer (10 mM Tris-HCl pH 8.0, 10 mM NaCl, 0.2% Igepal CA630, 33 μL Protease Inhibitor (Sigma, P8340)) on ice for 15 min, washed with 300 μL cold lysis buffer, and then incubated in 50uL of 0.5% SDS for 10min at 62°C. After heating, 170 μL of 1.47% Triton X-100 was added and incubated for 15min at 37°C. To digest chromatin 100U MboI and 25uL of 10X NEBuffer2 were added followed by overnight incubation at 37°C with agitation at 700rpm on a thermomixer. After incubation, MboI was inactivated by heating at 62°C for 20 minutes. The digested ends were filled and labeled with biotin by adding 37.5uL of 0.4mM biotin-14-dATP (Life Tech), 1.5 μL of 10mM dCTP, 10mM dTTP, 10mM dGTP, and 8uL of 5U/ul Klenow (New England Biolabs) and incubating at 23°C for 60 minutes with shaking at 500 rpm on a thermomixer. Then the samples were mixed with 1x T4 DNA ligase buffer (New England Biolabs), 0.83% Trition X-100, 0.1 mg/mL BSA, 2000U T4 DNA Ligase (New England Biolabs, M0202), and incubated for at 23°C for 4 hours with shaking at 300rpm on a thermomixer to ligate the ends. After the ligation reaction, samples were spun and pellets were resuspended in 550uL 10 mM Tris-HCl, pH 8.0. To digest the proteins and to reverse the crosslinks, 50 μL of 20mg/mL Proteinase K (New England Biolabs) and 57 μL of 10% SDS were mixed with the samples, and incubated at 55°C for 30 minutes, and then 67 μL of 5M NaCl were added followed by overnight incubation at 68°C. After cooling the samples, 0.8X Ampure (Beckman-Coulter) purification was performed. Next, the samples were sonicated to mean fragment length of 400 bp using Covaris M220 with the following parameters: 70 seconds duration at 10.0% duty factor, 50.0 peak power, 200 cycles per burst. To collect 200-600 bp size of fragmented DNA, two rounds of Ampure (Beckman-Coulter) beads purification was performed. The DNA labeled with biotin was purified using 100 μL of 10 mg/mL Dynabeads My One T1 Streptavidin beads (Invitrogen). The washed beads were transferred to the sample tube, incubated for 15 minutes at room temperature, and the supernatant was removed. Then the beads were washed twice by 600 μL of 1x Tween Wash Buffer with mixing for 2 minutes at 55°C. Then the beads were equilibrated once in 100 uL 1x NEB T4 DNA ligase buffer (New England Biolabs) followed by removal of the supernatant. To repair the fragmented ends and remove biotin from unligated ends, the beads were resuspended in 100uL of the following: 88 μL 1X NEB T4 DNA ligase buffer (New England Biolabs, B0202), 2 μL of 25mM dNTP mix, 5 μL of 10 U/μL T4 PNK (New England Biolabs), 4 μL of 3 U/μL NEB T4 DNA Polymerase (New England Biolabs), 1 μL of 5U/μL Klenow (New England Biolabs). The beads were incubated for 30 minutes at room temperature. The beads were washed twice by adding 600 μL of 1x Tween Wash Buffer, heating on a thermomixer for 2 minutes at 55°C with mixing. To add dA-tail, the beads were resuspended in 90 μL of 1X NEB Buffer2, 5 μL of 10mM dATP, and 5 μL of 5U/ul Klenow (exo-) (New England Biolabs). The beads were incubated for 30 minutes at 37°C. The beads were washed twice by adding 600 μL of 1x Tween Wash Buffer, heating on a thermomixer for 2 minutes at 55°C with mixing. Following the washes, the beads were equilibrated once in 100 μL 1x NEB Quick Ligation Reaction Buffer (New England Biolabs). Then the beads were resuspended again in 50 μL 1x NEB Quick Ligation Reaction Buffer. To ligate adapters, 2 μL of NEB DNA Quick Ligase (New England Biolabs) and 3 μL of Illumina Indexed adapter were added to the beads and incubated for 15 minutes at room temperature. The beads were washed twice with 600 μL of 1x Tween Wash Buffer, heating on a thermomixer for 2 minutes at 55°C with mixing. Then the beads were resuspended once in 100 μL 10 mM Tris-HCl, pH 8.0, followed by removal of the supernatant and resuspension again in 50 μL 10 mM Tris-HCl, pH 8.0. PCR amplification (8-9 cycles) was performed with 10 μL Fusion HF Buffer (New England Biolabs), 3.125 μL 10uM TruSeq Primer 1, 3.125 μL 10uM TruSeq Primer 2, 1 μL 10mM dNTPs, 0.5 μL Fusion HotStartII, 20.75 μL ddH20, 11.5 μL Bead-bound HiC library. Then PCR products underwent final purification using AMPure beads (Beckman-Coulter). Libraries were sequenced on Illumina HiSeq 4000.

### PLAC-seq library preparation

PLAC-seq experiments were performed as previously described^30^. Cells were crosslinked with 1% formaldehyde (w/v, methanol-free, ThermoFisher) for 15 minutes and quenched with 125mM glycine. The crosslinked pellets (2.5–3 million cells per sample) were incubated with 300ul of lysis buffer (10 mM Tris-HCl pH 8.0, 10 mM NaCl, 0.2% Igepal CA630, 33 μL, 1x complete protease inhibitor (Roche)) on ice for 15 min, washed with 500 μL cold lysis buffer, and then incubated in 50uL of 0.5% SDS for 10min at 62°C. After heating, 160 μL of 1.56% Triton X-100 was added and incubated for 15min at 37°C. To digest chromatin 100U MboI and 25uL of 10X NEBuffer2 were added followed by 2 hours incubation at 37°C with agitation at 900rpm on a thermomixer. After incubation, MboI was inactivated by heating at 62°C for 20 minutes. Digestion efficiency was confirmed by performing agarose gel electrophoresis of the samples. The digested ends were filled and labeled with biotin by adding 37.5uL of 0.4mM biotin-14-dATP (Life Tech), 1.5 μL of 10mM dCTP, 10mM dTTP, 10mM dGTP, and 8uL of 5U/ul Klenow (New England Biolabs) and incubating at 37°C for 60 minutes with shaking at 900 rpm on a thermomixer. Then the samples were mixed with 1x T4 DNA ligase buffer (New England Biolabs), 0.83% Trition X-100, 0.1 mg/mL BSA, 2000U T4 DNA Ligase (New England Biolabs, M0202), and incubated for at room temperature for 2 hours with shaking with slow rotation. The ligated cell pellets were resuspended in 125 ul of RIPA buffer with protease inhibitor and incubated on ice for 10 minutes. The cell lysates were sonicated using Covaris M220, and the sheared chromatins were spun at 14,000 rmp, 4°C for 15 minutes to clear the cell lysate. We saved 20 ul supernatant as input, and for the rest part, 100 ul of antibody-coupled beads were added to the supernatant sample, and then rotated in cold room at least 12 hours. For immunoprecipitation, 300 ul of M280 sheep anti-rabbit IgG beads (ThermoFisher) was washed with cold BSA/PBS (0.5 mg / mL bovine serum albumin in 1x phosphate buffered saline) for 4 times. After washing, 30 ug anti-H3K4me3 (Millipore, 04-745) with 1 mL cold BSA/PBS were added to the beads and incubated on a rotating platform at 4°C for at least 3 hours. After incubation, beads were washed with cold BSA/PBS, and resuspended in 600 ul RIPA buffer. The beads were washed with RIPA buffer (3 times), RIPA buffer + 0.16M NaCl (2 times), LiCl buffer (1 time), and TE buffer (2 times) at 4°C for 3 minutes at 1000 rpm. For reverse crosslinking, 163 ul extraction buffer (135 ul 1xTE, 15 ul 10% SDS, 12 ul 5M NaCl, 1 ul RnaseA (10mg/ml)) was added and incubated at 37°C for 1 hour at 1000 rpm, and 20 ug of proteinase K was added and incubated at 65°C for 2 hours at 1000rpm. After crosslinking, DNA was purified using Zymo DNA clean & concentrator and eluted with 50 ul of 10mM Tris (pH 8.0). For biotin enrichment, 25 ul of T1 Streptavidin Beads (Invitrogen) per sample were washed with 400 ul Tween wash buffer (5 mM Tris-HCl pH 8.0, 0.5 mM EDTA, 1 M NaCl, 0.05% Tween-20), and resuspended in 50 ul of 2x Binding buffer (10 mM Tris-HCl pH 7.5, 1 mM EDTA, 2 M NaCl). The purified 50 ul DNA sample was added to the 50 ul resuspended beads and incubated at room temperature for 15 minutes with rotation. The beads were washed with 500 ul of Tween wash buffer twice and washed with 100 ul Low EDTA TE (supplied by Swift Biosciences kit). Then beads were resuspended in 40 ul Low EDTA TE. Next, we used Swift Biosciences kit (Cat. No. 21024) for library construction with modified protocol as described below. The Repair I Reaction Mix was added to 40 ul sample beads and incubated at 37°C for 10 minutes at 800 rpm. The beads were washed with 500 ul Tween wash buffer twice and washed with 100 ul Low EDTA TE once. The Repair II Reaction Mix was added to the beads followed by incubation at 20°C for 20 minutes at 800 rpm. The beads were washed with 500 ul Tween wash buffer twice and washed with 100 ul Low EDTA TE once. Then 25 ul of the Ligation I Reaction Mix and Reagent Y2 was added to the beads followed by incubation at 25°C for 15 minutes at 800 rpm. The beads were washed with 500 ul Tween wash buffer twice and washed with 100 ul Low EDTA TE once. Then 50 ul of the Ligation II Reaction Mix was added to the beads followed by incubation at 40°C for 10 minutes at 800 rpm. The beads were washed with 500 ul Tween wash buffer twice and washed with 100 ul Low EDTA TE once, and resuspended in 21 ul 10mM Tris-HCl (pH 8.0). The amplification and purification were performed according to the Swift library kit protocols. Libraries were sequenced on Illumina HiSeq 4000.

### ChIP-seq data analysis

Each fastq file was mapped to mouse genome (mm10) with BWA^60^ -aln with “-q 5 -l 32 -k 2” options. PCR duplicates were removed using Picard MarkDuplicates (https://github.com/broadinstitute/picard) and the bigWig files were created using deepTools ^61^ with following parameters: bamCompare --binSize 10 --normalizeUsing RPKM --ratio subtract (or ratio). The deepTools was also used for generating heatmaps. Peaks were called with input control using MACS2 ^62^ with regular peak calling for narrow peaks (e.g. CTCF) and broad peak calling for broad peaks (e.g. H3K27me3, K3K4me1). Enhancer regions were characterized by the presence of both H3K4me1 peak and H3K27ac peak. We used DEseq2^63^ to calculate differences in peak levels between samples.

### RNA-seq data analysis

RNA-seq reads (paired-end, 100 bases) were aligned against the mouse mm10 genome assembly using STAR^64^. The mapped reads were counted using HTSeq^65^ and the output files from two replicates were subsequently analyzed by edgeR^66^ to estimate the transcript abundance and to detect the differentially expressed genes. Only genes that had H3K4me3 ChIP-seq peaks on TSS were used for downstream analysis (Fig. 1d, e). Differentially expressed genes were called by FDR < 0.01 and fold change > 2 thresholds. RPKM was calculated using an in-house pipeline.

### Hi-C data analysis

Hi-C reads (paired-end, 50 or 100 bases) were aligned against the mm10 genome using BWA^60^ -mem. Reads mapped to the same fragment were removed and PCR duplicate reads were removed using Picard MarkDuplicates. Raw contact matrices were constructed using in-house scripts with 10 or 40 kb resolution, and then normalized using HiCNorm^67^. We used juicebox pre^68^ to create hic file with -q 30 -f options. To visualize Hi-C data, we used Juicebox^68^ and 3D Genome Browser (http://www.3dgenome.org). Topological domain boundaries were identified at 40-kb or 10-kb resolution based on the directionality index (DI) score and a Hidden Markov Model as previously described^13^, and they were also identified based on insulation scores using peakdet (Billauer E, 2012. http://billauer.co.il/peakdet.html). The insulation score analysis was performed as previously described^69^ and insulation scores on TAD boundaries were calculated by taking the average value of scores that overlapped with TAD boundaries. The stripe calling was performed using a homemade pipeline (shared from Feng Yue lab, Penn State University) that is based on the algorithm proposed in a previous study^70^. We used HiCCUPS^68^ with options “-r 10000 -k KR -f 0.001 -p 2 -i 5 -d 50000” to identify Hi-C peaks as chromatin loops, and then we chose CTCF associated loops among them that were overlapped with convergently oriented CTCF ChIP-seq peaks in control cells. The aggregate analysis of CTCF associated loops were performed using APA^68^ with default parameters.

To assess global changes in TAD boundary strength between samples, we performed a comparison of each samples’ aggregated boundary contact profile (Supplementary Fig. 2c). First, to generate a consensus set of TAD boundaries we performed a simple merge between boundaries from clone 1 before auxin treatment (Clone 1, 0 hr) and boundaries from clone 2 before auxin treatment (Clone 2, 0 hr). Two filtering steps were used to generate the final set of consensus boundaries: 1) We discarded boundaries there were within 3.04 Mb of a chromosome start or end, because we would not be able to extract a submatrix of the correct size for the aggregate analysis; 2) We discarded boundaries > 200kb, because these often represent regions of disorganized chromatin between TADs, rather than true TAD boundaries. Next, we extracted a Hi-C sub-matrix for each boundary in each sample. Each sub-matrix consists of a window of 3.04 Mb centered on the midpoint of the boundary in question. These boundary sub-matrices were then averaged to generate one 3.04 Mb matrix representing the average boundary contact profile in a given sample. To facilitate comparison between samples, average boundary contact profiles were then normalized across samples using standard quantile normalization. We then made pairwise comparisons between samples by subtracting the average boundary contact profile of sample 1 from the average boundary contact profile of sample 2. The list of consensus TAD boundaries used here is the same as that described for the aggregate boundary analysis above.

### PLAC-seq data analysis

PLAC-seq reads (paired-end, 50 bases) were aligned against the mm10 genome using BWA^60^ -mem. Reads mapped to the same fragment were removed and PCR duplicate reads were removed using Picard MarkDuplicates. Filtered reads were binned at 10 kb size to generate the contact matrix. Individual bins that were overlapped with H3K4me3 peaks on TSSs were used for downstream differential contact analysis. For the peak calling, we used MAPS^71^ with default settings and FitHiChIP^72^ with coverage bias correction with default settings in 10 kb resolution.

For differential contact analysis, the raw contact counts in 10 kb resolution bins that have the same genomic distance were used as inputs for comparison. To minimize the bias from genomic distance, we stratified the inputs into every 10-kb genomic distance from 10 kb to 150 kb, and the other input bins with longer distances were stratified to have uniform size of input bins that were equal to that of 140–150 kb distance bins. Since each input showed negative binomial distribution, we used edgeR^66^ to get the initial set of differential interactions. We only used bins that have more than 20 contact counts in each sample of two replicates for downstream analysis. The significances of these differential interactions are either due to the difference in their H3K4me3 ChIP coverage or 3D contacts coverage. Therefore, the chromatin contacts overlapping with differential ChIP-seq peaks (FDR < 0.05, logFC < 0.5) were removed and only the chromatin contacts with the same level of H3K4me3 ChIP-seq peaks were processed. In this differential analysis, we used all bins for inputs that included non-significant interactions that were not identified by MAPS or FitHiChIP peak caller, because the majority of short-range interactions were not identified as significant peaks due to their high background and the changes in the short-range interactions might be also critical for gene regulation. We identified a large number of differentially changed short-range interactions even though many of them were not identified as significant peaks, and we observed a clear correlation between these differentially changed interactions and the changes of active enhancer levels on their anchor regions during neural differentiation, suggesting these interaction changes might reflect the biological changes. We used significance level with change direction (-/+ log_10_(p-value)) instead of fold change to show the changes of interactions, because fold change tends to be small value especially in short-range interactions even though the change is actually significant for biological aspects. To visualize the PLAC-seq contacts, we used WashU Epigenome Browser^73^.

### Active/inactive contact (AIC) ratio/value

The change of chromatin contacts on enhancer-promoter (E-P) is affected by the alteration of enhancer activities such as H3K27ac and H3K4me1 levels (Supplementary Fig. 5b, c), and it is also well known that gene expression levels have positive correlation with these active marks around their TSS. These findings suggest that information of contact counts itself should involve the information of enhancer activity. Furthermore, the majority of genes have multiple E-P contacts with variable changes of contact frequencies. Therefore, we designed AIC value to represent quantitative activity of multiple E-P contacts and aimed to show the relationship between gene regulation change and E-P contacts change without using any quantitative values of histone marks. First, we summed total contact counts on active elements and promoters in each gene. As for promoter-promoter (P-P) contacts, they have similar function as E-P contacts^74, 75^. However, it is still unclear that the same contact frequency of P-P contacts has the same effect as that of E-P contacts. Moreover, in H3K4me3 PLAC-seq datasets, P-P contacts correspond to peak-to-peak interactions that have generally higher contact counts than that of peak-to-non-peak interactions. Therefore, we divided the P-P contact counts by a certain integer that showed the highest correlation coefficient between gene expression changes and AIC value changes before summing total active contact counts. We tested integers from 0 to 10 to divide the P-P contact counts, and dividing by 3 and 8 showed the highest correlation coefficient between gene expression changes and AIC value changes in ESCs and NPCs, respectively. The simply summing of active contact counts is still not proper for comparison between different samples because they are affected by the difference of H3K4me3 peak levels on TSS in different samples. Therefore, in order to cancel the bias from the H3K4me3 peak levels in different samples, we also calculated total contact counts on inactive (non-active) regions and computed active/inactive contact (AIC) ratio on each gene by following formula.

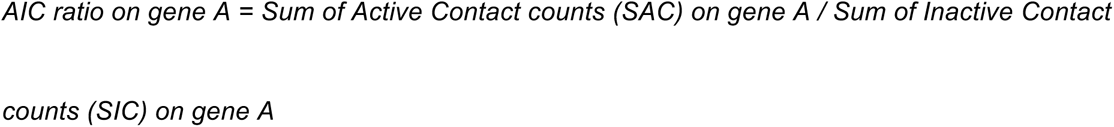

Next, we calculated the average of SICs from the comparing two samples on each gene, and multiplied them by AIC ratios to calculate AIC values. AIC values are computed as pseudo contact counts to perform differential analysis by edgeR^66^ after rounding them to their nearest integer. The bias from different H3K4me3 levels on TSS in different samples does not occur by multiplying the common average value of the SICs.

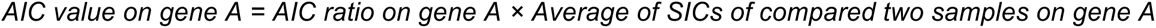

We also computed the changes of AIC values using Hi-C datasets in the same way, and we could observe comparable correlations with gene expression changes.

### Odds ratio calculation for CTCF-dependent E-P contacts enrichment

For Fig. 3a, b all PLAC-seq contacts (10 kb resolution) on promoter and enhancer were classified based on the distance from anchor sites (enhancer side or promoter side) to the nearest CTCF binding sites (Fig. 3a, categorized into 4×4 bins). They are also classified based on the number of CTCF motif sites around each anchor site (10 kb bin ±5 kb) (Fig. 3b, categorized into 5×5 bins). Then, we generated 2×2 tables based on whether they are CTCF-dependent contacts or not (FDR < 0.05) and whether they were categorized into the bin or not. Odds ratios and p-values on each 2×2 tables were calculated.

For Fig. 3c, d all PLAC-seq contacts (10 kb resolution) on promoter and enhancer were classified based on the distance from anchor sites (enhancer side or promoter side) to the nearest CTCF binding sites (Fig. 3c, categorized into 4×4 bins). They are also classified based on the number of CTCF motif sites around each anchor site (10 kb bin ±5 kb) (Fig. 3d, categorized into 5×5 bins). Then, we generated 2×2 tables based on whether they are on differentially down-regulated genes or not (FDR < 0.05) and whether they were categorized into the bin or not. For the E-P contacts on differentially down-regulated genes, chromatin contacts that were identified as significant by peak calling were counted (p value < 0.01). Odds ratios and p-values on each 2×2 tables were calculated.

For Fig. 4b, all genes were classified based on the distance to the nearest interacting enhancer and the number of enhancers around TSS (< 200 kb) (categorized into 3×3 bins). The distance to the nearest interacting enhancer is represented by the shortest genomic distance of significant PLAC-seq peaks on enhancers and promoters (p-value < 0.01). Then, we generated 2×2 tables based on whether they are CTCF-independent stably-regulated genes or not (FDR < 0.05) and whether they were categorized into the bin or not. Odds ratios and p-values on each 2×2 tables were calculated. In Fig. 4c and Supplementary Fig. 10f, the same analysis as Fig. 4b was performed in CTCF-dependent down-regulated genes and CTCF-dependent up-regulated genes.

### 3D modelling

We used the Strings & Binders Switch (SBS) polymer model^39, 76^ to dissect the 3D spatial organization of the *Vcan* gene region in wild type and CTCF depleted NPC cells. In the SBS view, a chromatin filament is modelled as a Self-Avoiding Walk (SAW) chain of beads, comprising different specific binding sites for diffusing cognate molecular factors, called binders. Different types of binding sites are visually represented by distinct colors. Beads and binders of the same color interact with an attractive potential, so driving the folding of the chain. All binders also interact unspecifically with all the beads of the polymer by a weaker energy affinity (see below). We estimated the optimal number of distinct binding site types describing the locus and their arrangement along the polymer chain by using the PRISMR algorithm, a previously described machine learning based procedure^40^. In brief, PRISMR takes as input a pairwise experimental contact map (e.g. Hi-C) of the studied genomic region and, via a standard Simulated Annealing Monte Carlo optimization, returns the minimal number of different binding site types and their arrangement along the SBS polymer chain, which best reproduce the input contact map. Next, we ran Molecular Dynamics (MD) simulations of the inferred SBS polymers so to produce a thermodynamics ensemble of single molecule 3D conformations.

We focused on the genomic region chr13:89,200,000-92,000,000 (mm10) encompassing the mouse *Vcan* gene, in wild type and CTCF depleted NPC cells. Applied to our Hi-C contacts data of the region, at 10kb resolution, the PRISMR algorithm^40^ returned in both cases polymer models made of 6 different types of binding sites. In our simulations, beads and binders interact via standard potentials of classical polymer physics studies^77^ and the system Brownian dynamics is defined by the Langevin equation. By using the LAMMPS software^78^, we ran massive parallel MD simulations so producing an ensemble of, at least, 10^2^ independent conformations. We started our MD simulations from initial SAW configurations and let the polymers evolve up to 10^8^ MD time steps when the equilibrium globule phase is reached. We explored a range of specific and non-specific binding energies in the weak biochemical energy range, respectively from 3.0K_B_T to 8.0K_B_T and from 0K_B_T to 2.7K_B_T, where K_B_ is the Boltzmann constant and T is the system temperature. For the sake of simplicity, those affinity strengths are the same for all the different types. All details about the model and MD simulations are described in ^39, 40^. To better highlight the locations of the *Vcan* gene and its regulatory elements in the two different cases, we produced a coarse-grained version of the polymers. We interpolated the coordinates of the beads with a smooth third-order polynomial curve and used the POV-ray software (Persistence of Vision Raytracer Pty. Ltd) to produce the 3D images.

For the model derived contact maps, we computed the average contact frequencies from our MD derived ensemble of 3D polymer model conformations for each cell type. We followed a standard approach that considers a pair of polymer sites in contact if their physical distance is lower than a threshold distance^40^. To compare model contact maps with corresponding Hi-C data in each cell type, we used the HiCRep stratum adjusted correlation coefficient (SCC)^79^, a bias-corrected correlation designed for Hi-C comparison, with a smoothing parameter h=5 and an upper bound of interaction distance equal to 1.5Mb. To compute the model frequencies of multiple contacts, we proceeded similarly. Specifically, fixed a viewpoint site k, we accounted for a triple contact (i,j,k) between k and any pair of sites i,j along the locus if their relative physical distances were all lower than the threshold distance.

### CTCF motif deletion and tethering dCas9-CTCF

The CRISPR/Cas9 system was used to delete CTCF motif nearby *Vcan* promoter. The sequences of the DNA targeted are listed below (the protospacer adjacent motif is underlined). The guide RNAs were generated using GeneArt Precision gRNA Synthesis Kit (Invitrogen).

5’-TTCAGCACAAGCGGAAAATA<u>GGG</u>-3’,

5’-CTGCTTGCAGTTGGGTGTTT<u>CGG</u>-3’

Transfection of gRNA and Cas9 ribonucleoprotein (EnGen SpyCas9, New England Biolabs) into mESCs was performed using Neon Transfection System, 10 ul tip kit (Life Technologies). The cells were grown for approximately one week, and individual colonies were picked into a 96-well plate. After expanding cells, genotyping by PCR and Sanger sequencing were performed to confirm the motif deletion.

For the generation of dCas9-CTCF tethered cell lines, plasmids containing sequences for dCas9 and CTCF were generated by modifying lenti-dCas-VP64-Blast (a gift from Feng Zhang, Addgene #61425). The VP64 cassette was replaced by CTCF sequences to generate dCas9-CTCF and neomycin resistant marker that was taken from pAC95-pmax-dCas9VP160-2A-neo (a gift from Rudolf Jaenisch, Addgene 48227) was inserted. To generate gRNA plasmids to recruit dCas9, the gRNA oligos were inserted into the backbone vector (pSPgRNA, a gift from Charles Gersbach, Addgene Plasmid #47108). The gRNA was designed to target the top and bottom strand of *Vcan* promoter-proximal region which is close to the deleted CTCF motif locus. The sequences of the DNA targeted are listed below (the protospacer adjacent motif is underlined).

5’-CCTGCCTCCTTGGACAGAGA<u>CGG</u>-3’ (for top strand)

5’-GTCCCTTCCGTCTCTGTCCA<u>AGG</u>-3’ (for bottom strand)

The plasmids for dCas9-CTCF and gRNA were extracted using PureLink HiPure Plasmid Midiprep Kit (Invitrogen). For the electroporation, 350 ng of dCas9-CTCF plasmids and 600 ng of gRNA plasmids (1 ul) were added to 0.1–0.2 million mESCs resuspended in 10 ul Buffer R (Invitrogen), and electric pulse was delivered with the setting of 1200 V, 20 ms, and 2 pulses. After culturing approximately 10 days, individual colonies were picked and genotyping and western blotting were performed to confirm the sequences from the transfection plasmids and their protein expression. For deletion of enhancer region that is interacting with *Vcan* promoter. The sequences of the DNA targeted are listed below (the protospacer adjacent motif is underlined).

5’-AGGAACGGCCCATTCCCGAG<u>GGG</u>-3’,

5’-CAATCAATAATAACACGCAT<u>AGG</u>-3’

Generating gRNA and transfection of gRNA and Cas9 ribonucleoprotein into mESCs were performed in the same way as the deletion of CTCF motif was done. Genotyping by PCR and Sanger sequencing were performed to confirm the deletion.

### Analysis of CTCF-occupied promoter (COP) genes in multiple mouse tissues

To analyze the CTCF ChIP-seq signals around promoters, we calculated fold changes of sample RPKM over input RPKM in each 50-bp bin and summed them in each promoter region (TSS ±10 kb) when the 50-bp bins were located at the regions of optimal IDR thresholded ChIP-seq peaks. Then correlation coefficient between these summed CTCF ChIP-seq signals and RNA-seq RPKM values across 9 mouse tissues was computed in each gene. Heatmap was generated for genes with high correlation coefficient (> 0.6). The values in the heatmap were calculated by log_2_(summed ChIP-seq signals / average value of all tissues) for promoter-proximal CTCF signal and log_2_(RPKM / average RPKM of all tissues) for gene expression. Lineage specificity of transcription was measured by Shannon entropy^43^. For DNA methylation levels around promoters, DNA methylation rates at CBSs (motif sequences ±100 bp) were calculated and averaged in each promoter region (TSS ±10 kb). PhastCons score^80^ was used for conservation analysis. The highest pahstCons score at each CTCF motif locus was represented as the conservation score of each CBS.

### Data accessibility

All datasets generated in this study have been deposited to Gene Expression Omnibus (GEO), with accession number GSE94452. Hi-C dataset analyzed in Supplementary Fig. 2f–h was provided from Dr. Benoit Bruneau^28^. All mouse tissue datasets in Fig. 5 and Supplementary Fig. 12 were downloaded from the ENCODE portal^81^ (https://www.encodeproject.org/); accessions for datasets are described in Supplementary Table 6.

## Acknowledgments

We thank Drs. Elphrege Nora and Benoit Bruneau for exchanging datasets and reagents. We would like to give special thanks to Samantha Kuan for operating the sequencing instruments and Tristin Liu and Zhen Ye for helping experiments. We would like to acknowledge the help of Drs. Victor Lobanenkov and Arshad Desai for giving helpful advice and the help of Drs. Feng Yue, Xiaotao Wang, Ivan Juric, and Armen Abnousi for sharing computational pipelines. We would also like to give special thanks to Drs. Rongxin Fang, Yanxiao Zhang, Ramya Raviram, Anthony Schmitt, and Sora Chee for sharing helpful information and protocols, as well as all the other members of the Ren laboratory. This work was supported by the Ludwig Institute for Cancer Research (B.R.), NIH (1U54DK107977-01) (B.R.), NIH (1U54DK107965) (H.Z.), a Ruth L. Kirschstein Institutional National Research Award from the National Institute for General Medical Sciences, T32 GM008666 (J.D.H.), and a Postdoc fellowship from the TOYOBO Biotechnology Foundation (N.K.).

## Author Information

The authors declare no competing financial interests. Correspondence and requests for materials should be addressed to B.R.

## Author contributions

N.K., H.I., and B.R. conceived the project. N.K., H.I., X.X., and H.Z. engineered cell lines. N.K., R.H., J.D.H., and Z.Y. carried out library preparation. N.K. and F.M. performed cell cycle analysis. N.K., S.B., M.C., M.N., D.G., and B.L. performed data analysis. M.Y., M.H., and J.D. contributed to computational analysis and experimental design. N.K. and B.R. wrote the manuscript. All authors edited the manuscript.

---

**Supplementary Fig. 1.**
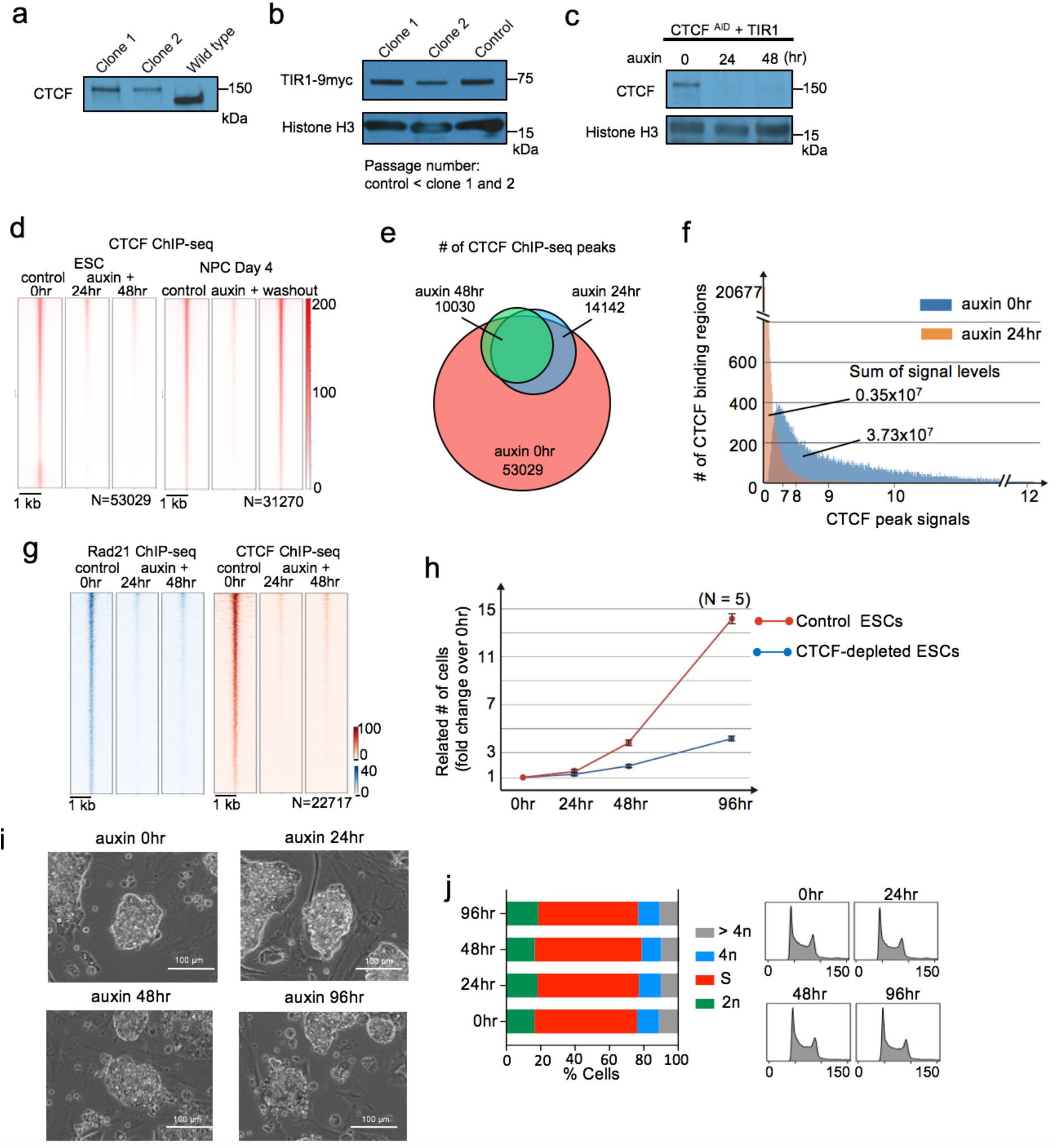
Depletion of CTCF and characterization of CTCF-depleted cells. **a, b,** Western blot showing AID-tagged CTCF and wild type CTCF (a) and the expression of TIR1 protein in 2 clones of mESCs. The TIR1 expression in these clones that went through multiple passages were comparable to that in control cells that have much smaller passage number. **c**, Western blot showing the acute depletion of CTCF protein after 24 and 48 hours of auxin treatment. **d,** Heatmaps showing CTCF ChIP-seq signals centered at all regions of CTCF peaks identified in the control cells and CTCF occupancy at the same regions in CTCF-depleted cells at each time point of ESC and NPC stages. The CTCF peaks disappeared almost completely after auxin treatment and recovered after washing out auxin. **e,** Venn-diagram comparing the number of CTCF ChIP-seq peaks identified in control and CTCF-depleted ESCs at each time point. **f,** Histogram showing the number of CTCF binding regions in *y*-axis and their CTCF ChIP-seq signal levels in *x*-axis. The CTCF signal levels in control cells and auxin treated cells were calculated on the CTCF peak regions identified in the control cells. **g,** Heatmaps comparing the Rad21 ChIP-seq signals centered at all regions of Rad21 peaks identified in control ESCs and in CTCF-depleted ESCs at each time point (left, blue heat map). The CTCF occupancy at the same regions in the same samples are also shown (right, red heat map). **h,** Growth curves of mouse ESCs with or without auxin treatment. **i,** Bright-field microscopy images of mouse ESC colonies before and after multiple days of auxin treatment. **j,** Cell cycle analysis by flow cytometry using propidium iodide staining in control and 24, 48, and 96 hours auxin treated CTCF-depleted ESCs suggests that CTCF-depleted cells can pass through cell cycle checkpoints.

**Supplementary Fig. 2.**
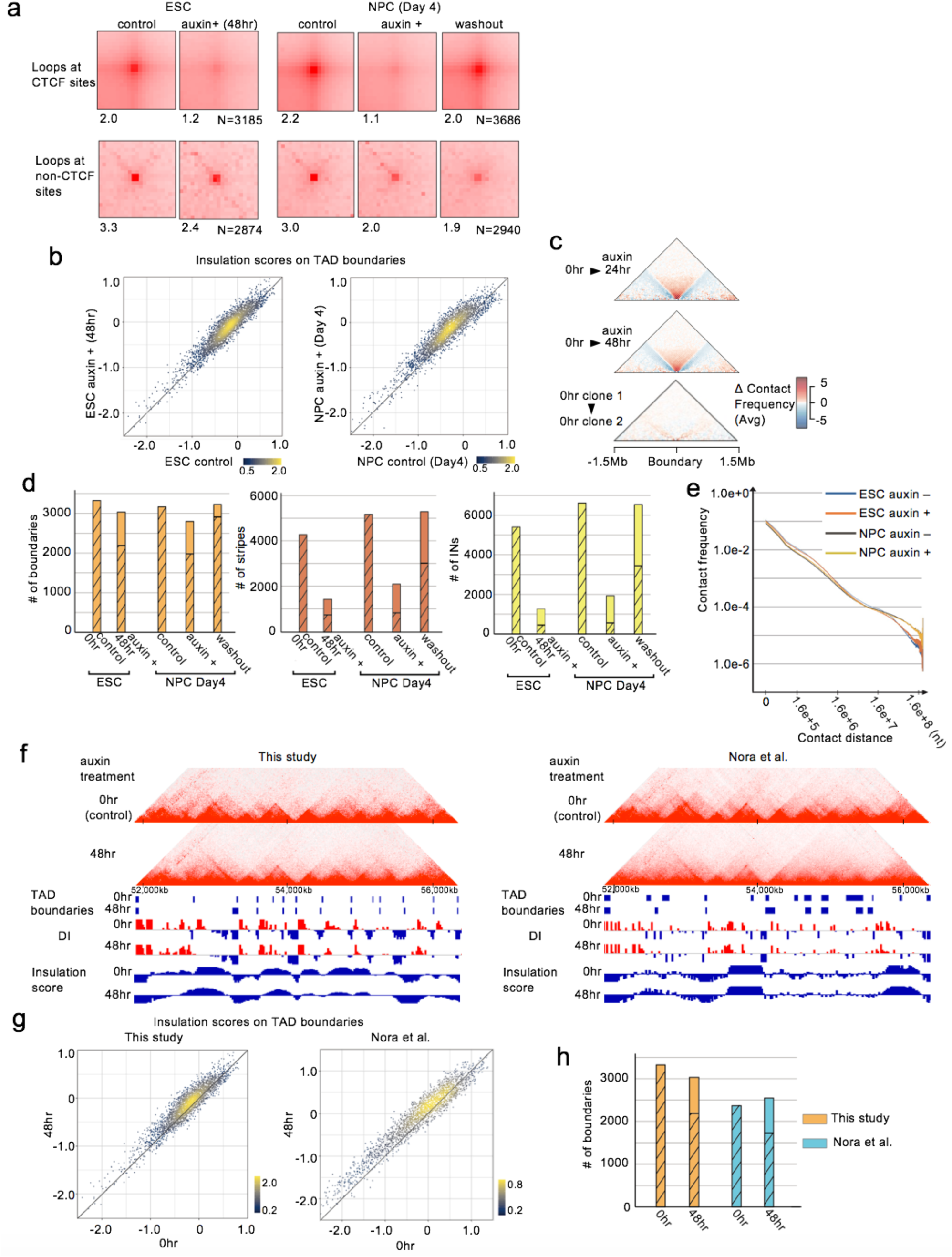
Severe disruption of chromatin architecture upon CTCF loss. **a,**Scatter plots showing insulation scores at TAD boundaries. A higher score denotes lower insulation. Increased insulation scores in varying degrees were observed in CTCF-depleted ESCs and NPCs. **b,** Aggregate boundary analysis showing the average change in boundary strength between samples. Each triangle is a contact map showing the difference in the average contact profile at TAD boundaries between two time points. The bottom column shows that there is little difference in the average boundary profile between the two control samples. **c,** The number of TAD boundaries (left), stripes (middle), and insulated neighborhoods (INs) in control, CTCF-depleted, and auxin washout cells. The hatched bars indicate the number of boundaries/stripes/INs that overlapped with boundaries/stripes/INs identified in control cells. **d,** Hi-C contact frequencies at each genomic distance are plotted. Overall profiles of Hi-C contact frequencies were not affected by the loss of CTCF but affected by cell differentiation. **e,** APA (aggregate peak analysis) on Hi-C peak loci on convergent CTCF binding sites identified in control ESCs and NPCs (n=3185 (ESCs), 3686 (NPCs), > 100-kb looping range). APA was performed in control and CTCF-depleted cells in ESC and NPC stages and also performed after washing out auxin in NPCs. APA on Hi-C peak loci that have no CTCF binding sites (n=2874 (ESCs), 2940 (NPCs), > 100-kb looping range) was also performed. The scores on the bottom represent the focal enrichment of peak pixel against pixels in its lower left. **f–h,** Hi-C datasets generated in this study and Nora et al. study^28^. Genome browser snapshots showing Hi-C contact heatmaps, TAD boundaries, directionality indices (DIs), and insulation scores analyzed in the two independent studies at the same genomic region in control and CTCF-depleted cells (f). Scatter plots showing insulation scores at all TAD boundaries (g). The number of TAD boundaries in control and CTCF-depleted cells from the two studies. The hatched bars indicate the number of boundaries that overlapped with the boundaries identified in control cells (h). The degree of weakening of TAD boundaries was comparable between the two studies.

**Supplementary Fig. 3.**
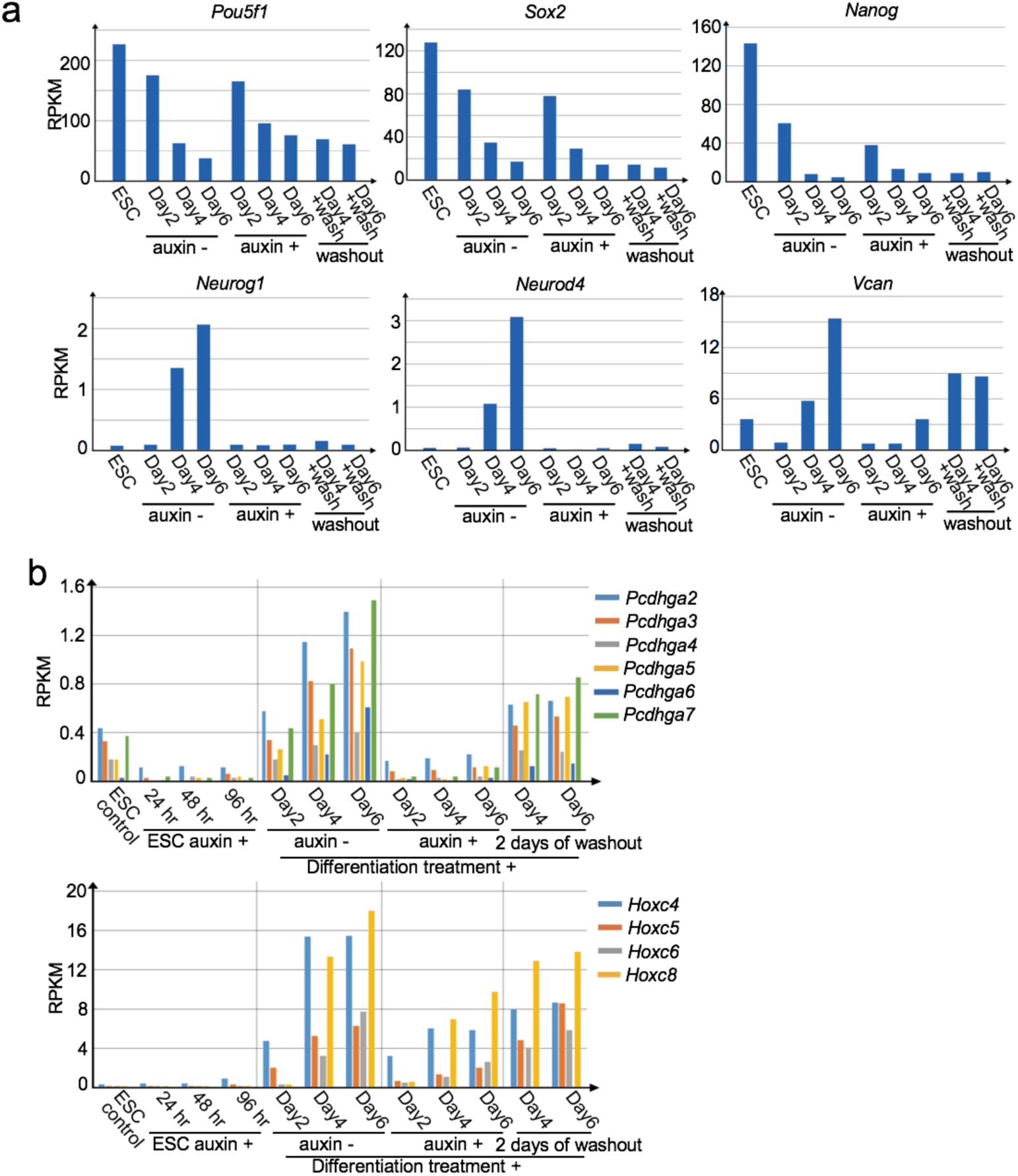
Examples of gene expression changes during neural differentiation in control and CTCF-depleted cells. **a,**Gene expression profiles of pluripotent marker genes (*Pou5f1*, *Sox2*, *Nanog*) and examples of induction failure gene upon CTCF loss that is important for nervous system development (*Neurog1, Neurod4, Vcan*) in control and CTCF-depleted cells during differentiation from ESC to NPC and 2 days after washing out auxin in NPCs. **b,** Gene expression profiles of *Pcdhga* and *Hoxc* gene clusters during multiple days of auxin treatment in ESCs and during differentiation from ESC to NPC in control and CTCF-depleted cells followed by washing out auxin in NPCs

**Supplementary Fig. 4.**
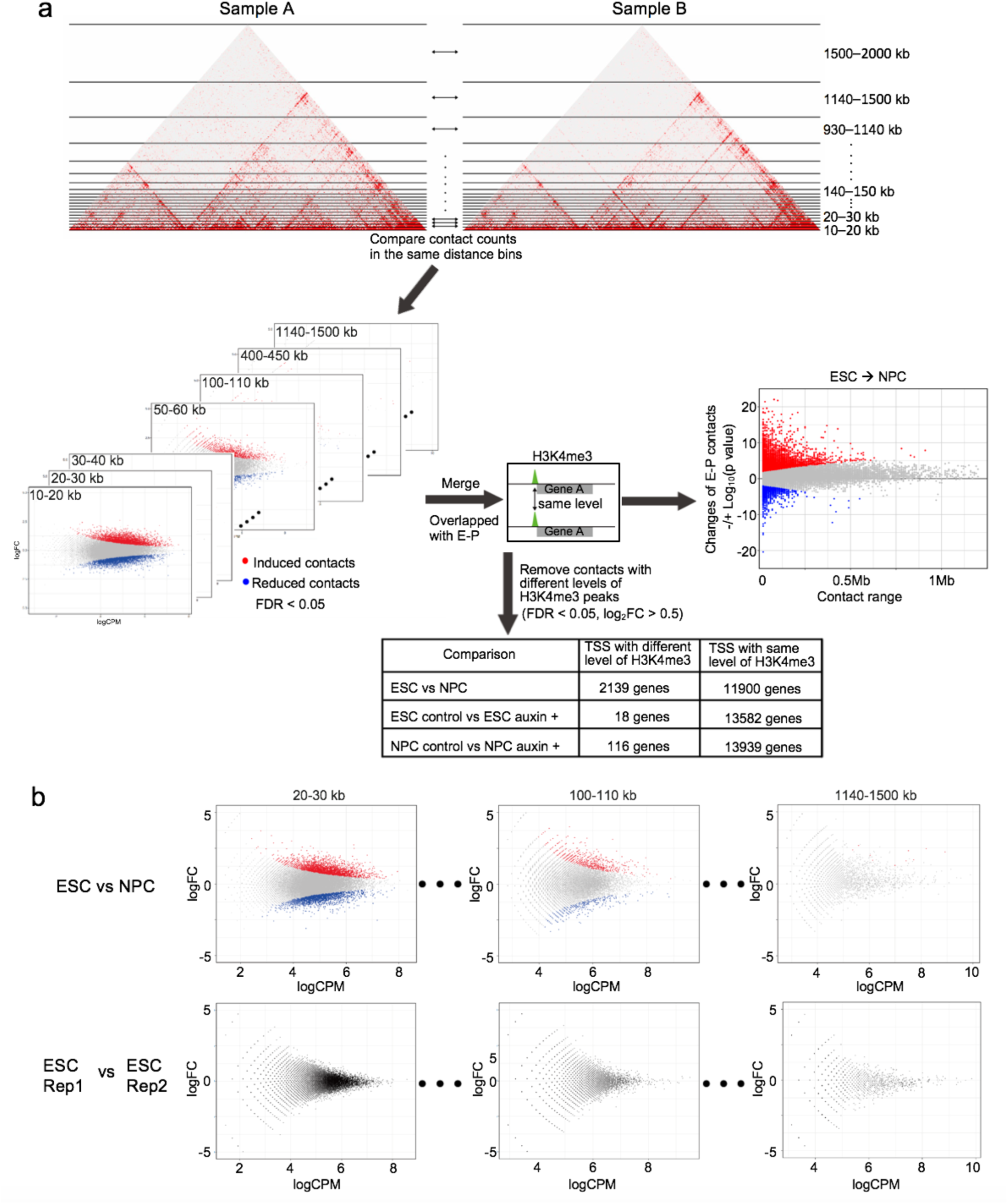
Differential chromatin interaction analysis of PLAC-seq datasets. **a,**Schematic representation of differential interaction analysis in H3K4me3 PLAC-seq datasets. The mapped contact counts in 10-kb resolution bins that have the same genomic distance were compared separately. The input contact matrix bins were stratified into every 10-kb genomic distance from 10-kb to 150-kb and the other bins with longer distances were stratified to have a uniform number of input bins that were equal to that of 140–150 kb distance bins. Two sets of these discrete inputs of two biological replicates were compared using a negative binomial model, edgeR^58^. The interactions anchored at the differential H3K4me3 ChIP-seq peaks between the conditions (FDR < 0.05, |logFC| > 0.5) were removed and only the genes with the same level of H3K4me3 ChIP-seq peaks on promoters were processed in downstream analysis (see Methods). **b,** Differential interaction analysis between ESC and NPC (top) in each distance-stratified 10-kb interval. The difference between ESC replicate 1 and replicate 2 are also shown for comparison (bottom).

**Supplementary Fig. 5.**
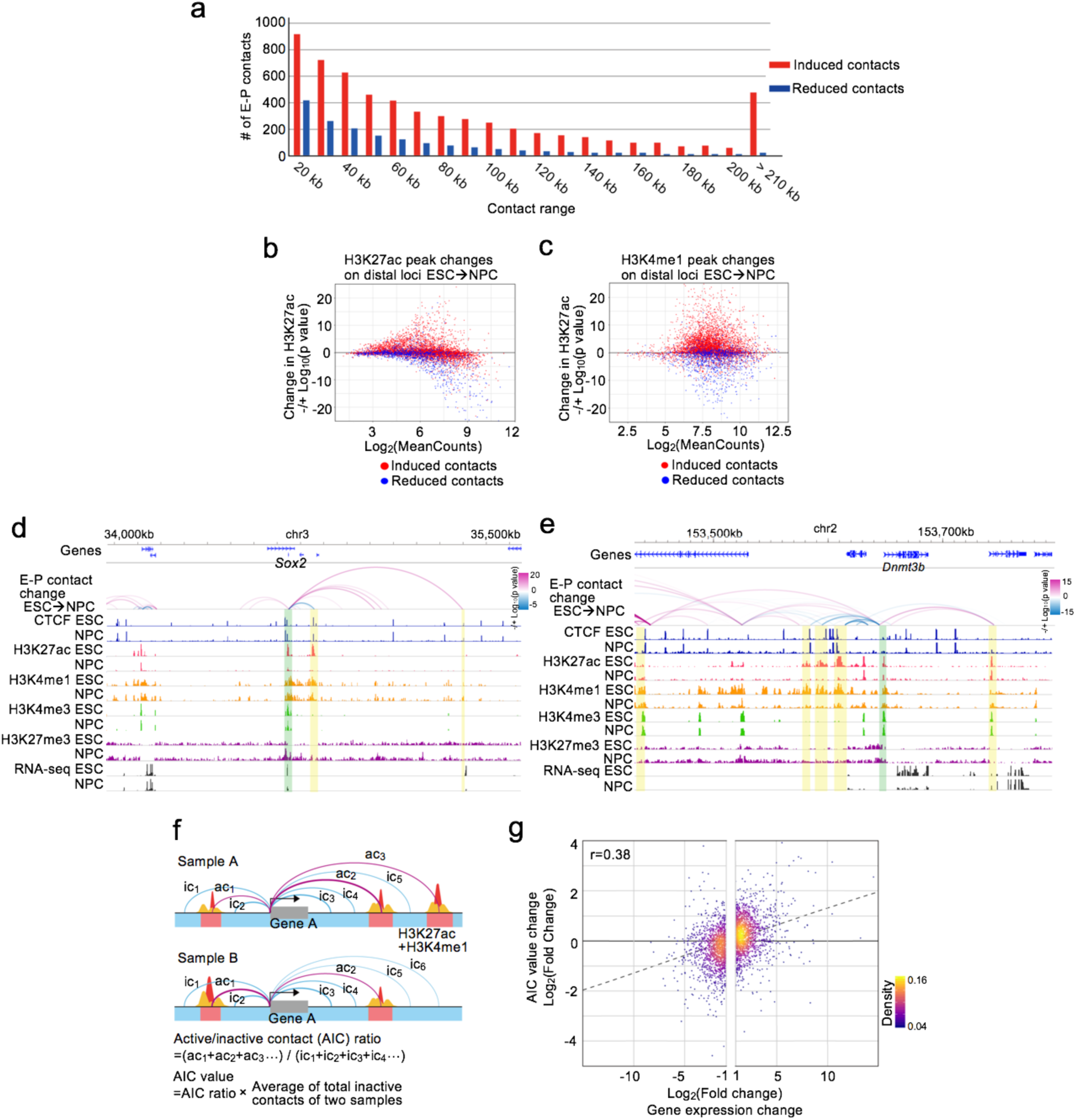
Changes of enhancer-promoter contacts during neural differentiation were associated with the changes of chromatin modification and gene expression. **a,**Histogram showing the number of differentially changed enhancer-promoter (E-P) contacts between control ESCs and NPCs and their genomic distances. While the majority of them (53%) were less than 50 kb distance, a large number of long-range E-P contacts were induced during differentiation. Red bars: significantly induced E-P contacts. Blue bars: significantly reduced E-P contacts. **b, c,** Scatter plots showing changes of ChIP-seq peak levels of H3K27ac (c) and H3K4me1 (d) on distal element loci of induced (red) and reduced (blue) E-P contacts during neural differentiation. **d, e,** Genome browser snapshots of regions around *Sox2* gene (a) and *Dnmt3b* gene (b) that were down-regulated during neural differentiation. The arcs show the changes of chromatin contacts on active elements and promoters identified in differential interaction analysis between ESCs and NPCs. The colors of arcs represent degrees of interaction change between the conditions (blue to red, -/+log_10_(p-value)). The promoter regions of these genes and interacting enhancer regions are shown in green and yellow shadows, respectively. CTCF, H3K4me1, H3K27ac, H3K4me3, H3K27me3 ChIP-seq and RNA-seq in ESCs and NPCs (day 4) are also shown. **f,** Schematic representation of Active/Inactive contact (AIC) model to see the correlation between changes of multiple E-P contacts and gene expression levels. H3K27ac and H3K4me1 peaks are shown as red and yellow peaks, respectively, and regions where these two types of peaks are overlapping are defined as active elements (red colored regions). Promoter-centered chromatin contacts on these active elements are shown as red arcs (active contacts) and the other chromatin contacts are shown as blue arcs (inactive contacts). AIC ratio/value was calculated by the formulae described on the bottom (see Methods). **g,** Scatter plots showing the changes of AIC values (y-axis) and gene expression levels (x-axis) in differentially expressed genes during neural differentiation with linear approximation.

**Supplementary Fig. 6.**
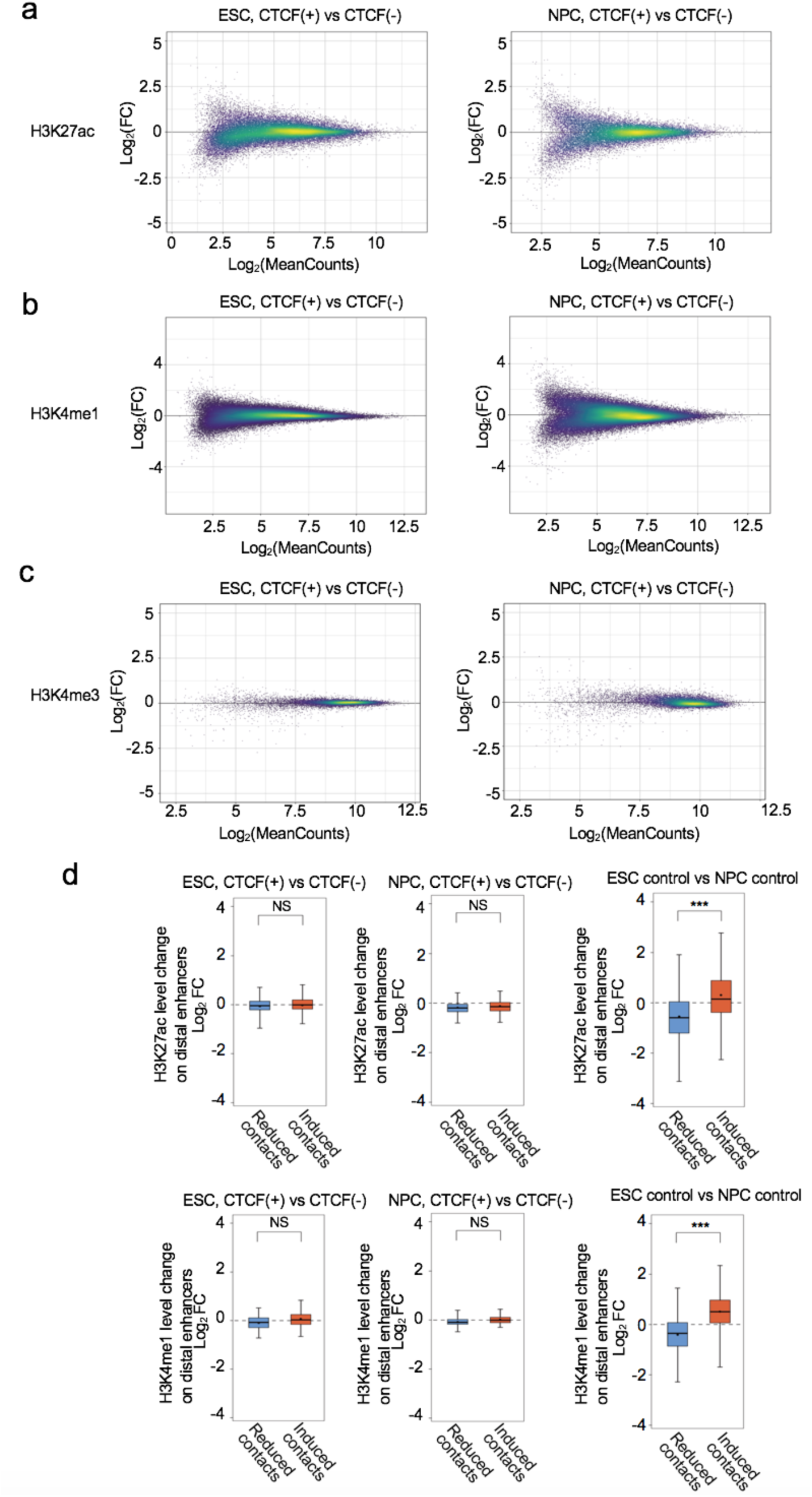
CTCF loss does not alter histone modification landscape. **a–c,** Scatter plots showing the changes of H3K27ac (a) and H3K4me1 (b) ChIP-seq signal levels upon CTCF loss on all significant peak regions in ESCs (left) and NPCs (right). The changes of H3K4me3 ChIP-seq signal levels on all peak regions on transcription start sites (TSSs) are also shown (c). **d,** Boxplots showing the changes of H3K27ac (top) and H3K4me1 (bottom) ChIP-seq signal levels on distal element loci of all analyzed E-P contacts. The changes upon CTCF depletion in ESC (left) and NPC stage (middle) and the changes during neural differentiation (right) are shown. (NS not significant, *** p value < 0.001, two-tailed t-test).

**Supplementary Fig. 7.**
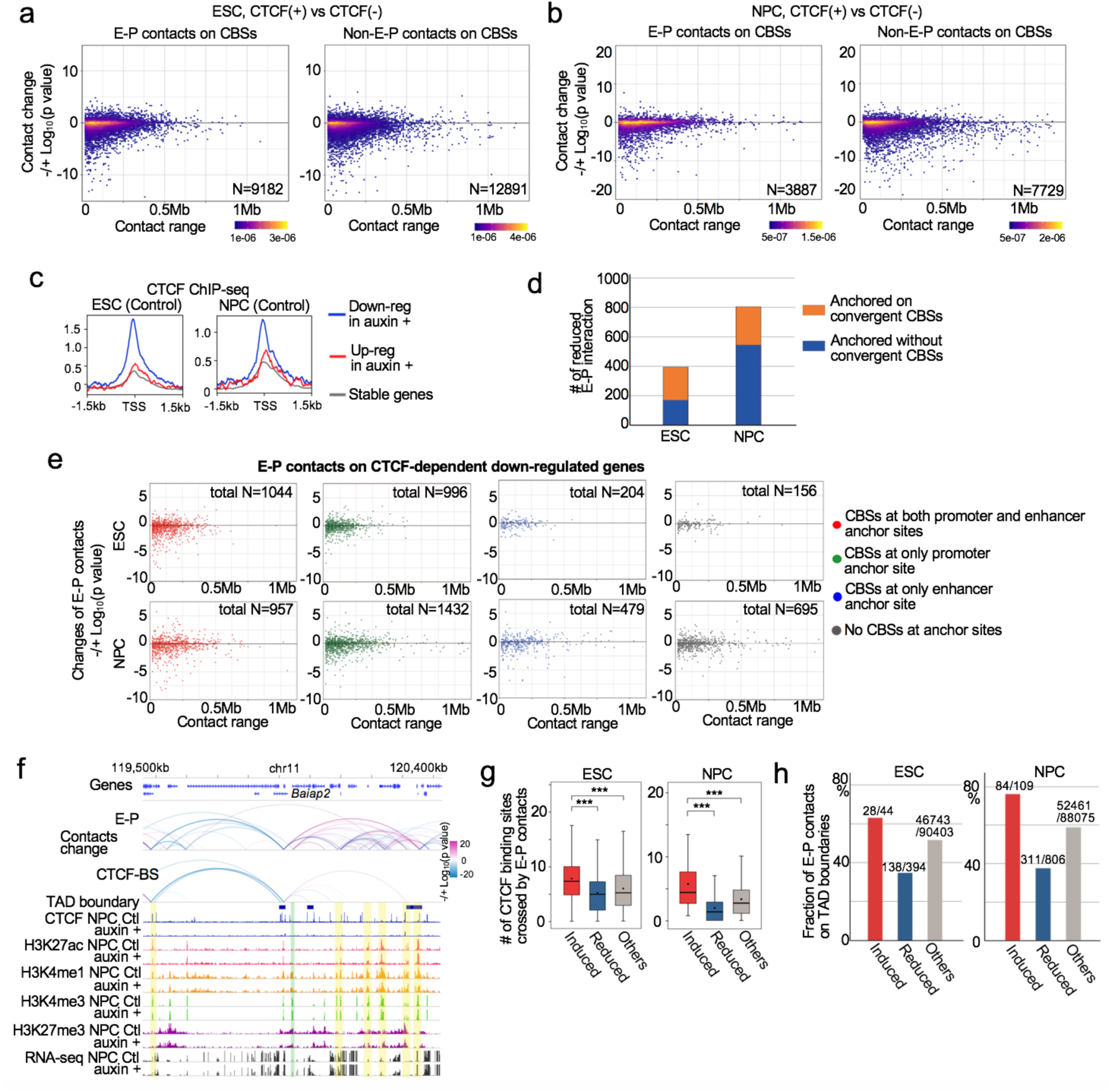
Changes of chromatin contacts upon CTCF loss and features of CTCF-dependent enhancer-promoter contacts. **a, b,** Scatter plots showing changes of H3K4me3 PLAC-seq contacts (*y*-axis) on convergently oriented CTCF binding sites and their loop ranges (*x*-axis). Chromatin contacts in CTCF-depleted cells were compared to the chromatin contacts in control cells in ESC (a) and NPC stage (day 4) (b). The plots were classified based on whether they are on promoters and enhancers (E-P) (left) or not (Non-E-P) (right). **c,** Average enrichments of CTCF ChIP-seq signals on TSSs of CTCF-dependent up-regulated genes (red), CTCF-dependent down-regulated genes (blue), and the other CTCF-independent stably regulated genes (gray) upon CTCF loss in ESC (left) and NPC stage (right). **d,** The number of CTCF-dependent differentially reduced E-P contacts anchored on convergently oriented CBSs (orange) and the CTCF-dependent E-P contacts anchored without convergently oriented CBSs (blue) in ESCs (left) and NPCs (right). **e,** Scatter plots showing changes of chromatin contacts anchored on promoters of CTCF-dependent down-regulated genes and distal active elements (*y*-axis). Genomic distances between their two loop anchor sites are plotted in *x*-axis. Chromatin contacts in CTCF-depleted cells were compared to the chromatin contacts in control cells in ESC (top) and NPC stage (day 4) (bottom). The E-P contacts were classified based on following four conditions. CBSs are located at both anchor sites (10 kb bin ±5 kb) (red dots). CBSs are located only at promoter side of their anchor sites (green dots). CBSs are located only at distal element side of their anchor sites (blue dots). CBSs are located at neither anchor sites (gray dots). The total number of analyzed bin pairs are added. **f,** Genome browser snapshots of a region around *Baiap2* gene up-regulated upon CTCF loss in NPCs. The arcs show the changes of chromatin contacts on enhancers and promoters (E-P) and chromatin contacts on CTCF binding sites (CTCF-BS). The colors of arcs represent degrees of interaction change from control cells to CTCF-depleted cells (blue to red, -/+log_10_(p-value)). The promoter regions of *Baiap2* gene and interacting enhancer regions are shown in green and yellow shadows respectively. Newly formed E-P contacts were observed in *Baiap2* gene promoter. CTCF, H3K4me1, H3K27ac, H3K4me3, H3K27me3 ChIP-seq, and RNA-seq in control and CTCF-depleted NPCs, and TAD boundaries in control cells are also shown. **g,** Boxplots showing the number of CTCF binding sites that were located in between the two anchor sites of the following three types of E-P contacts. Significantly induced E-P contacts upon CTCF loss (red), significantly reduced E-P contacts upon CTCF loss (CTCF-dependent) (blue), and unchanged E-P contacts upon CTCF loss (CTCF-independent) (gray). (*** p value < 0.001, two-tailed t-test). **h,** The fraction of E-P contacts that were overlapped with TAD boundaries in ESC (left) and NPC stage (left). Red: significantly induced E-P contacts upon CTCF loss. Blue: significantly reduced E-P contacts upon CTCF loss (CTCF-dependent). Gray: unchanged E-P contacts upon CTCF loss (CTCF-independent).

**Supplementary Fig. 8.**
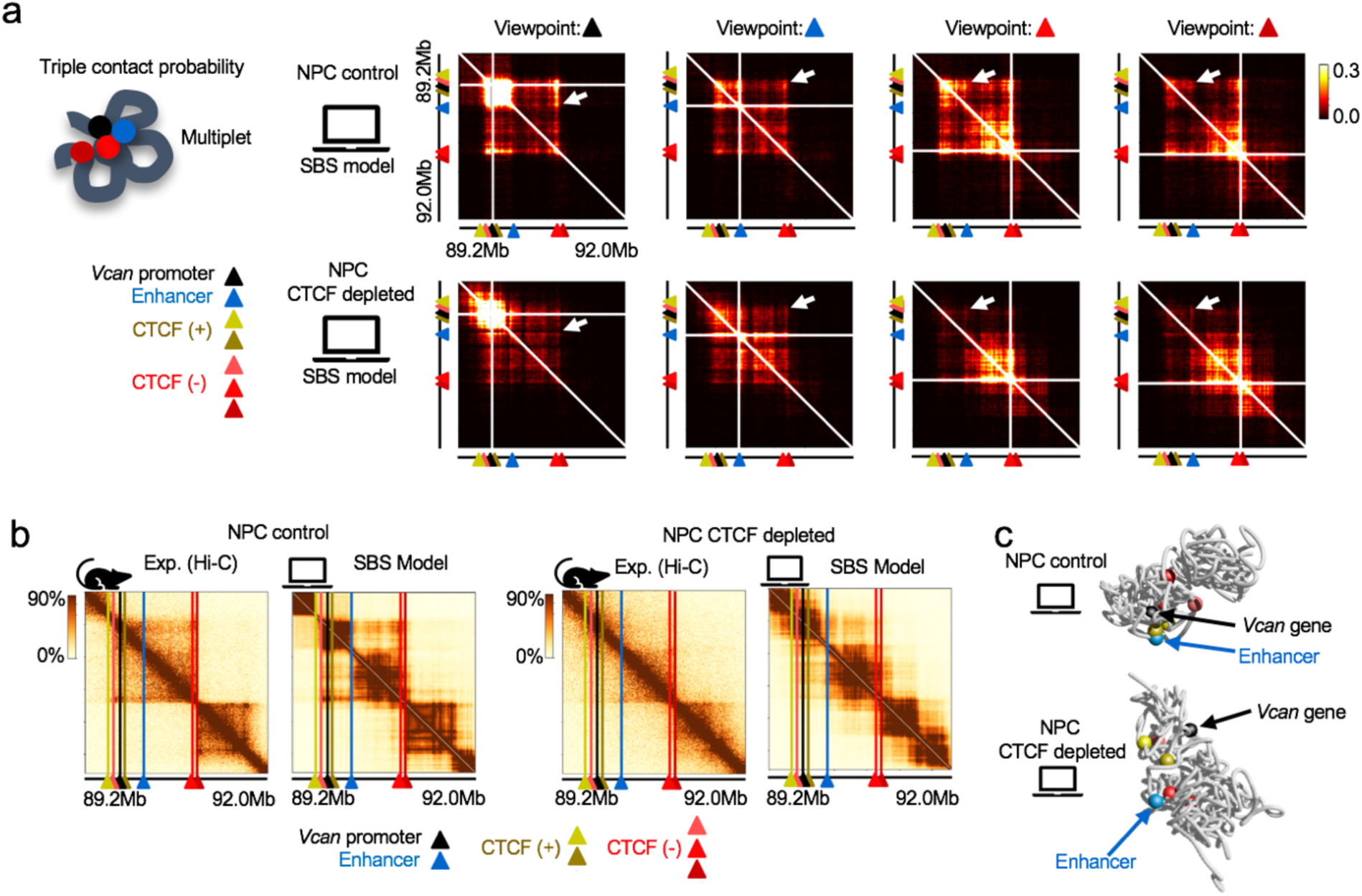
Verification of chromatin contacts change upon CTCF loss by SBS polymer model. **a,**The Strings & Binders Switch (SBS) model showing triplet interactions between *Vcan* promoter (black), distal enhancer (blue) and distal CTCFs (-) (reds), that weaken upon CTCF depletion (white arrows). Heatmaps from each viewpoint in control NPCs and CTCF-depleted NPCs are shown. CTCFs (+) (browns) and CTCFs (-) (reds) are convergently oriented. **b,** Hi-C contact maps (left) of the *Vcan* locus in NPC control and auxin treated cells are well reproduced by the SBS polymer model (right) (HiCRep stratum adjusted correlation SCC = 0.76 and SCC = 0.62 respectively). Genomic positions of *Vcan* promoter (black), distal enhancer (blue) and relevant motif-oriented CTCF sites (brown and red) are shown by colored triangles. **c,** SBS derived 3D structures of the control NPCs and CTCF-depleted NPCs, with relevant elements indicated by colored beads (the color code is the same as panel b). The models capture the loss of contact between *Vcan* promoter and its distal enhancer upon CTCF depletion.

**Supplementary Fig. 9.**
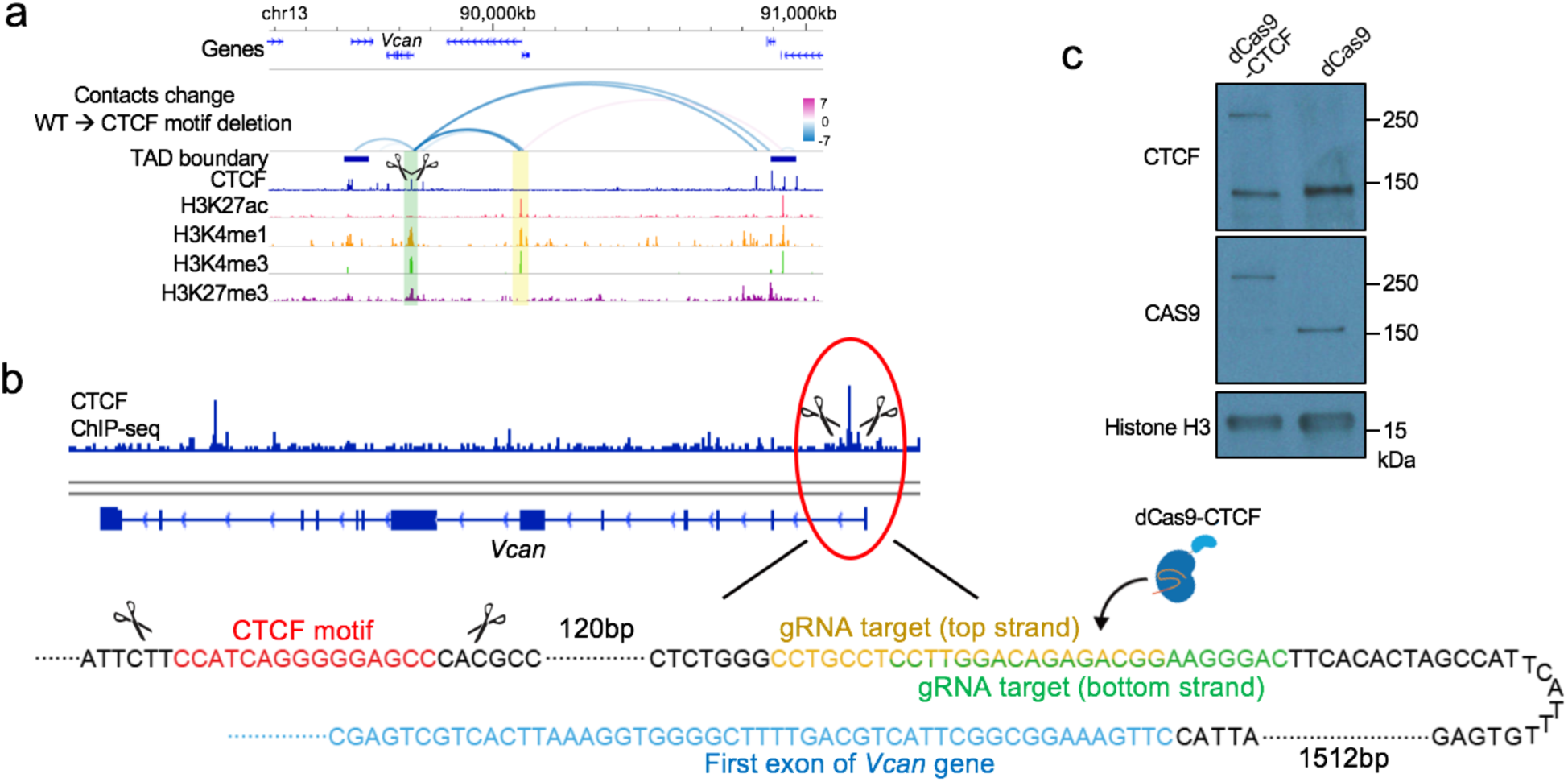
Rescue experiments using dCas9-CTCF. **a,**Genome browser snapshots of a region around *Vcan* gene. The arcs show the changes of chromatin contacts anchored on *Vcan* promoter, distal enhancer, and CTCF binding sites identified in differential interaction analysis between wild type NPCs and the promoter-proximal CTCF motif sequences deleted NPCs. The colors of arcs represent degrees of interaction change upon the deletion of CTCF motif sequences (blue to red, -/+log_10_(p-value)). The promoter region and interacting enhancer region are shown in green and yellow shadows, respectively. CTCF, H3K27ac, H3K4me1, H3K4me3, and H3K27me3 ChIP-seq, and TAD boundaries in wild type NPCs are also shown. The deletion of the single CTCF binding site leaded to disruption of its E-P contact. **b,** Schematic representation of the dCas9-CTCF rescue experiments showing targeted DNA sequences for CTCF motif deletion and dCas9-CTCF recruitment. **c,** Western blot showing that the cells transfected with dCas9-CTCF or dCas9 alone were successfully expressing dCas9-CTCF or dCas9 proteins.

**Supplementary Fig. 10.**
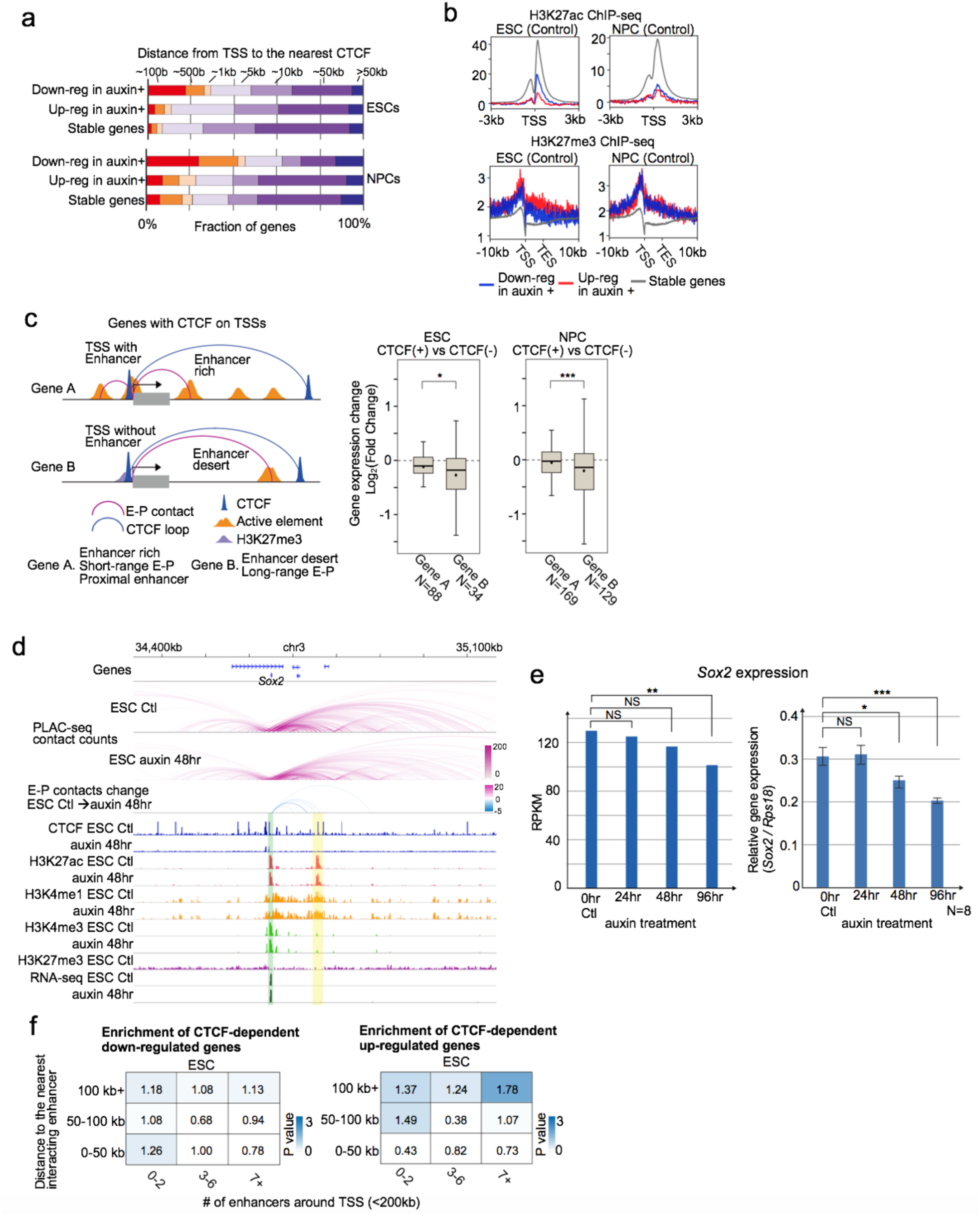
Mechanisms of CTCF-dependent/-independent gene regulation. **a,**Fraction of genes classified based on genomic distance from TSS to the nearest CTCF ChIP-seq peak. CTCF-dependent down-regulated genes, up-regulated genes, and CTCF-independent stably regulated genes in ESCs (top) and NPCs (bottom) are shown. **b,** Average enrichments of H3K27ac ChIP-seq signals on TSSs of CTCF-dependent up-regulated genes (red), CTCF-dependent down-regulated genes (blue), and the other CTCF-independent stably regulated genes (gray) in ESC (left) and NPC stage (right). Average enrichments of H3K27me3 ChIP-seq signals on TSSs and transcription end sites (TESs) are also shown (bottom). **c,** Schematic representation showing two types of genes that had CTCF bindings on their TSSs (< 1 kb). In Gene A, the shortest E-P contact (PLAC-seq peak signal p-value < 0.01) is shorter than 50 kb genomic distance and there are 7 enhancers or more around TSS (< 200 kb). In Gene B, the shortest E-P contact is longer than 50 kb and there are 2 enhancers or less around TSS. Boxplots showing gene expression changes upon CTCF loss in Gene A and Gene B in ESCs (left) and NPCs (right), suggesting that down-regulation upon loss of promoter-proximal CTCF is milder when genes are surrounded by proximal enhancers and regulated by short-range E-P contacts. (* p value < 0.05 and *** p value < 0.001, two-tailed t-test) **d, e,** Genome browser snapshot of a region around *Sox2* gene (d) whose expression level was not changed significantly 24 or 48 hours after CTCF depletion in ESCs in RNA-seq and qPCR (e). The arcs show PLAC-seq contact counts in control (top) and CTCF-depleted ESCs (middle) at every 10-kb bin. The changes of chromatin contacts on enhancers and *Sox2* gene promoter identified in differential interaction analysis between the two conditions are also shown (bottom). *Sox2* gene promoter and interacting super enhancer are shown in green and yellow shadows, respectively. CTCF, H3K4me1, H3K27ac, H3K4me3, and H3K27me3 ChIP-seq, RNA-seq in control and CTCF-depleted ESCs are shown. Consistent with the relatively stable gene regulation upon CTCF loss, the changes of E-P contacts were also mild even though convergent CTCF binding sites are located around its anchor sites. **f,** Enrichment analysis of CTCF-dependent down-regulated genes (left) and up-regulated genes (right) categorized based on the distance to the nearest interacting enhancer (vertical columns) and the number of enhancers around TSS (< 200 kb) (horizontal columns) in ESCs. Enrichment values are shown by odds ratio (scores in boxes) and p-values (color). The distance to the nearest interacting enhancer is represented by the shortest genomic distance of significant PLAC-seq peaks on enhancers and promoters (p-value < 0.01). (see Fig. 4 c for the same analysis in NPCs and Methods).

**Supplementary Fig. 11.**
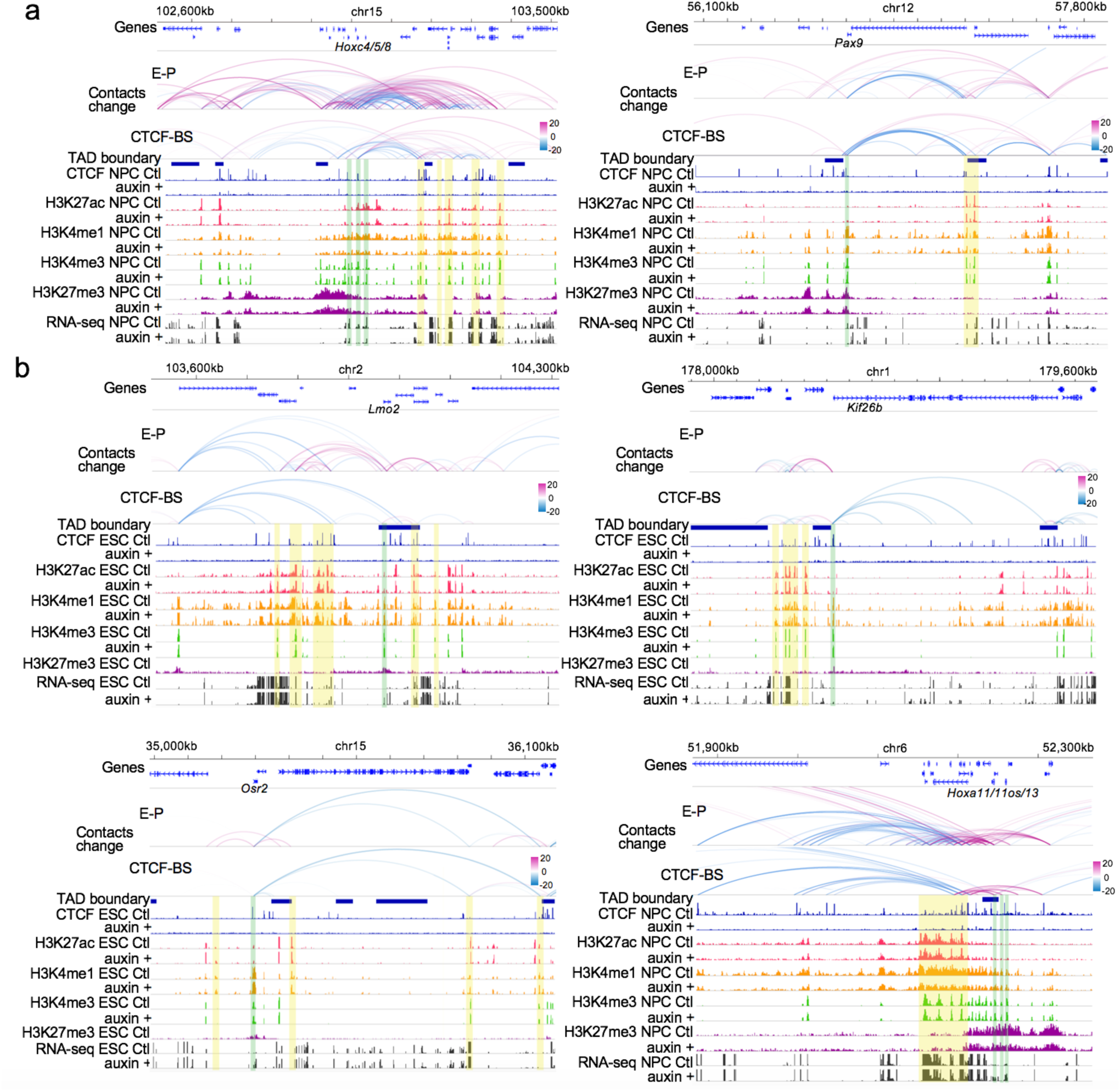
Examples of changes in enhancer-promoter contacts upon CTCF loss. **a,**Genome browser snapshots of regions around *Hoxc4, Hoxc5, Hoxc8* genes and *Pax9* gene that were down-regulated upon CTCF loss in NPCs. Data display shown here is the same as Supplementary Fig. 7f. **b,** Genome browser snapshots of regions around *Lmo2, Kif26b, Osr2, Hoxa11, and Hoxa13* genes that were up-regulated upon CTCF loss in ESCs or NPCs. Data display shown here is the same as Supplementary Fig. 7f.

**Supplementary Fig. 12.**
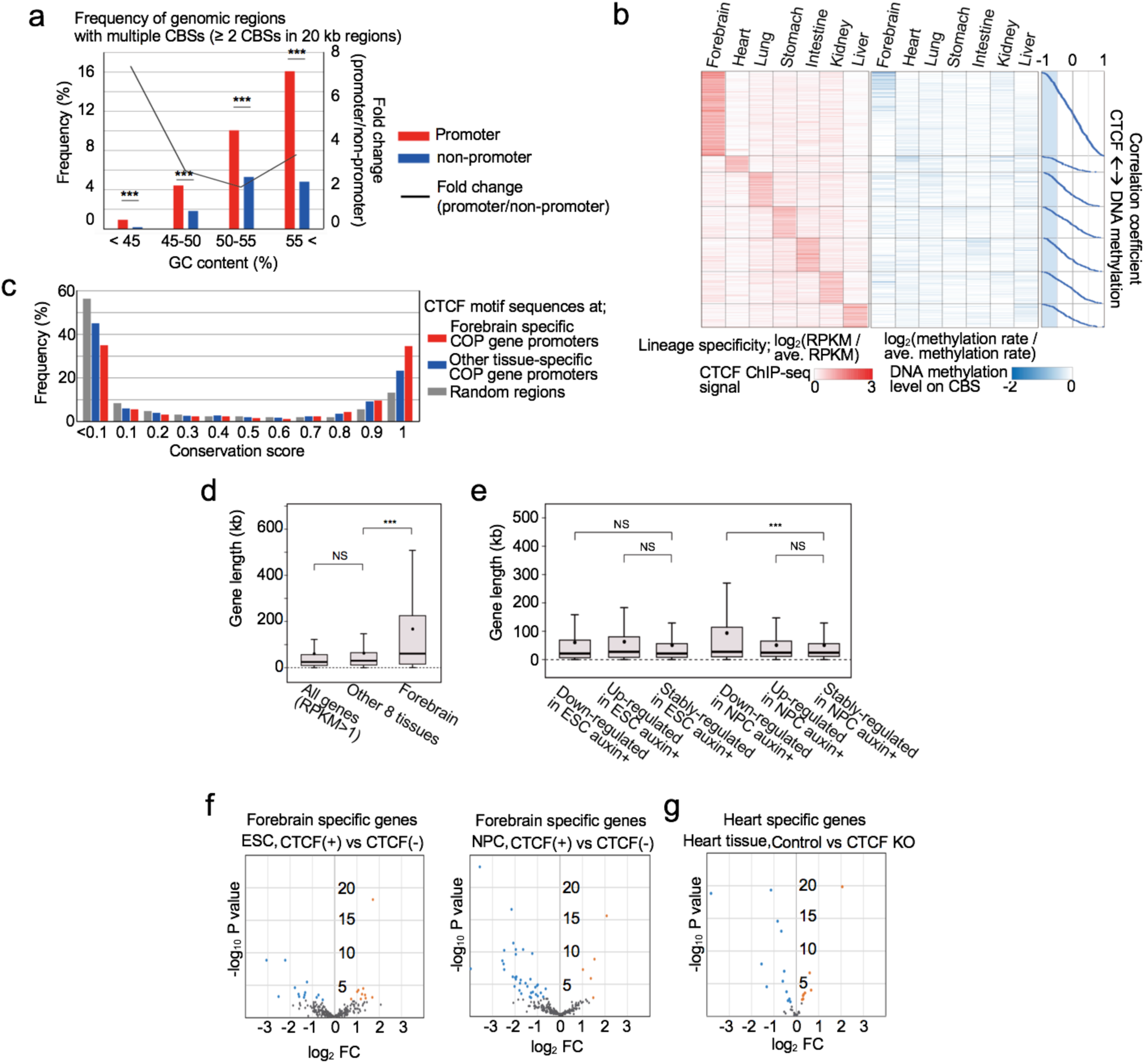
Features of tissue-specific CTCF occupied promoter genes. **a,**Histogram showing frequencies of genomic regions that has 2 CBSs or more in all analyzed 9 tissues. They are classified based on GC content levels. Red bars show the frequencies in promoter regions (TSS ±10 kb) and blue bars show the frequencies in random regions other than promoter regions. Black line shows fold change between the two groups in each GC content level. (*** p value < 0.001, two-tailed t-test). **b,** Heatmap showing lineage-specificity of DNA methylation levels at CBSs (motif sequences ±100 bp) in promoter regions of genes that are shown in Fig. 5c. The DNA methylation levels at multiple CBSs in the same promoter region (TSS ±10 kb) were averaged. The lineage-specificity of DNA methylation levels shown in the heatmap are calculated by log_2_(DNA methylation level / average methylation level of all tissues), and heatmap was sorted by correlation coefficient between CTCF ChIP-seq signal levels and the DNA methylation levels across the multiple tissues in each tissue group. Each correlation coefficient is shown in the scatter plots (right), indicating that only a partial of CTCF bindings had negative correlation with DNA methylation levels (r < −0.5, highlighted in blue). **c,** Histogram showing frequencies of CTCF motif sequences and their PhastCons conservation scores. Red and Blue bars show CTCF motif sequences at forebrain-specific CTCF occupied gene promoters and other tissue-specific CTCF occupied gene promoters, respectively. Gray bars show CTCF motif sequences at random regions. **d,** Boxplots showing gene length of lineage-specific CTCF occupied promoter genes that had high correlation coefficient (> 0.6) shown in Fig 5b, c. Forebrain-specific genes and the rest of the other lineage-specific genes are shown at right and middle, respectively. All genes whose RNA-seq RPKM value is more than 1 in at least one tissue sample were computed as control (left). (NS not significant, *** p value < 0.001, two-tailed t-test) **e,** Boxplots showing gene length of CTCF-dependent down-regulated genes, CTCF-dependent up-regulated genes, and CTCF-independent stably-regulated genes in ESCs and NPCs. (NS not significant, *** p value < 0.001, two-tailed t-test). **f,** Volcano plots showing the gene expression changes of the forebrain-specific CTCF-occupied genes between control cells and CTCF-depleted cells in ESCs (left) and NPCs (right). The larger number of forebrain-specific genes were down-regulated in NPCs upon CTCF loss. **g,** Volcano plots showing gene expression changes of the heart-tissue-specific CTCF-occupied genes between control heart tissue and CTCF knockout heart tissue.

**Supplementary Table 1.**

List of NGS sample information.

**Supplementary Table 2.**

List of CTCF ChIP-seq peaks in control and CTCF-depleted cells.

**Supplementary Table 3.**

List of TAD boundaries and Stripes in control and CTCF-depleted cells.

**Supplementary Table 4.**

Gene expression changes upon CTCF loss in each time point during neural differentiation.

**Supplementary Table 5.**

List of differentially changed enhancer-promoter (promoter-promoter) interactions.

**Supplementary Table 6.**

List of ENCODE datasets that were used for Fig. 5 and Supplementary Fig. 12.

