## Supplementary material for "CTCF Promotes Long-range Enhancer-promoter Interactions and Lineage-specific Gene Expression in Mammalian Cells": Table S6

**Supplementary Table 6; ENCODE datasets in Fig. 5 and Extended Data Fig.11**

| **Experiment** | **Data type** | **Tissue** | **Laboratory** |
| --- | --- | --- | --- |
| ENCSR677HXC | CTCF ChIP-seq | Forebrain, Postnatal 0 days | Richard Myers, HAIB |
| ENCSR491NUM | CTCF ChIP-seq | Heart, Postnatal 0 days | Richard Myers, HAIB |
| ENCSR002ZAG | CTCF ChIP-seq | Intestine, Postnatal 0 days | Richard Myers, HAIB |
| ENCSR143WOK | CTCF ChIP-seq | Kidney, Postnatal 0 days | Richard Myers, HAIB |
| ENCSR041SMK | CTCF ChIP-seq | Liver, Postnatal 0 days | Richard Myers, HAIB |
| ENCSR418SBY | CTCF ChIP-seq | Lung, Postnatal 0 days | Richard Myers, HAIB |
| ENCSR104QEN | CTCF ChIP-seq | Stomach, Postnatal 0 days | Richard Myers, HAIB |
| ENCSR000CDZ | CTCF ChIP-seq | Thymus,  male adult 8 weeks | Bing Ren, UCSD |
| ENCSR000CFJ | CTCF ChIP-seq | Bone marrow macrophage, male adult 8 weeks | Bing Ren, UCSD |
| ENCSR362AIZ | RNA-seq | Forebrain, Postnatal 0 days | Barbara Wold, Caltech |
| ENCSR526SEX | RNA-seq | Heart, Postnatal 0 days | Barbara Wold, Caltech |
| ENCSR331XCE | RNA-seq | Intestine, Postnatal 0 days | Barbara Wold, Caltech |
| ENCSR173PJN | RNA-seq | Kidney, Postnatal 0 days | Barbara Wold, Caltech |
| ENCSR096STK | RNA-seq | Liver, Postnatal 0 days | Barbara Wold, Caltech |
| ENCSR982MRY | RNA-seq | Lung, Postnatal 0 days | Barbara Wold, Caltech |
| ENCSR178GUS | RNA-seq | Stomach, Postnatal 0 days | Barbara Wold, Caltech |
| ENCSR000BYV | RNA-seq | Thymus,  Male adult 8 weeks | Thomas Gingeras, CSHL |
| ENCSR000CHO | RNA-seq | Bone marrow macrophage, Male adult 8 weeks | Bing Ren, UCSD |
| ENCSR020GGM | WGBS | Forebrain, Postnatal 0 days | Joe Ecker, Salk |
| ENCSR397YEG | WGBS | Heart, Postnatal 0 days | Joe Ecker, Salk |
| ENCSR353IFP | WGBS | Intestine, Postnatal 0 days | Joe Ecker, Salk |
| ENCSR128HOP | WGBS | Kidney, Postnatal 0 days | Joe Ecker, Salk |
| ENCSR550CYA | WGBS | Liver, Postnatal 0 days | Joe Ecker, Salk |
| ENCSR409HKJ | WGBS | Lung, Postnatal 0 days | Joe Ecker, Salk |
| ENCSR286OOJ | WGBS | Stomach, Postnatal 0 days | Joe Ecker, Salk |
